# Quantifying constraint in the human mitochondrial genome

**DOI:** 10.1101/2022.12.16.520778

**Authors:** Nicole J. Lake, Wei Liu, Stephanie L. Battle, Kristen M. Laricchia, Grace Tiao, Daniela Puiu, Alison G. Compton, Shannon Cowie, John Christodoulou, David R. Thorburn, Hongyu Zhao, Dan E. Arking, Shamil R. Sunyaev, Monkol Lek

## Abstract

Mitochondrial DNA (mtDNA) has an important, yet often overlooked, role in health and disease. Constraint models quantify the removal of deleterious variation from the population by selection, representing a powerful tool for identifying genetic variation underlying human phenotypes^1–4^. However, a constraint model for the mtDNA has not been developed, due to its unique features. Here we describe the development of a mitochondrial constraint model and its application to the Genome Aggregation Database (gnomAD), a large-scale population dataset reporting mtDNA variation across 56,434 humans^5^. Our results demonstrate strong depletion of expected variation, suggesting most deleterious mtDNA variants remain undiscovered. To aid their identification, we compute constraint metrics for every mitochondrial protein, tRNA, and rRNA gene, revealing a spectrum of intolerance to variation. We characterize the most constrained regions within genes via regional constraint, and positions across the entire mtDNA via local constraint, showing their enrichment in pathogenic variation and functionally critical sites, including topological clustering in 3D protein and RNA structures. Notably, we identify constraint at often overlooked sites, such as rRNAs and non-coding regions. Lastly, we demonstrate how these metrics can improve the discovery of mtDNA variation underlying rare and common human phenotypes.

## Main

Mitochondria produce the majority of cellular energy supplies and play a key role in many other cellular processes including signaling pathways, redox homeostasis, cell fate decisions, immune response, and regulation of metabolism^6–8^. Mitochondria originate from bacteria acquired by eukaryotic cells, and their prokaryotic origin is reflected by the circular mitochondrial genome (mtDNA), which is maintained and expressed in the mitochondria separately from the nuclear genome^9^. The human mtDNA is ∼16.5 kb in length and encodes for 13 proteins, as well as 22 transfer RNAs (tRNAs) and two ribosomal RNAs (rRNA) required for their translation^10,11^. These proteins encode subunits of complexes I, III, IV and V in the oxidative phosphorylation (OXPHOS) pathway, which are enzymes central to energy generation and cellular metabolism. The mtDNA has several unique features which differentiate it from the nuclear genome. This includes maternal inheritance, absence of introns, multiple copies per cell (e.g. 100s-1000s), germline bottleneck, distinct mutational mechanisms, and a higher rate of mutation and fixation relative to the nuclear genome^7,12^. The percentage of mtDNA molecules with a variant, known as the heteroplasmy level, can therefore range from 0-100%. Variants are heteroplasmic when they are in a fraction of mtDNA copies, or homoplasmic when found in all.

Variants in the mtDNA can cause mitochondrial and cellular dysfunction, and accordingly have been linked to many human phenotypes. This includes causing rare mitochondrial diseases, which are clinically heterogeneous disorders causing ‘*any symptom, in any organ, at any age*’^11^, as well as increasing the risk of common diseases such as autism^13^, cancer^14^, Alzheimer’s and Parkinson’s diseases^15^. Mitochondrial variation has also been linked to traits including glucose and insulin levels^16^, height^17^, and aging^18^. Despite its importance in health and disease, the effect of most mtDNA variants remains unknown. The classification of mtDNA variants is particularly challenging owing to the unique features of this genome, but also due to the paucity of tools for predicting mtDNA variant effect^12^. For example, zero *in silico* pathogenicity predictors exist for rRNA or non-coding mtDNA variants, and only one missense predictor is recommended in ACMG/AMP mtDNA guidelines^12^. Another tool that has not been available for mtDNA is a constraint model, which has been used to quantify the removal of deleterious nuclear genome variation from the population by selection^1^. This method compares the observed level of variation in a population to that expected under neutrality, as calculated by a mutational model. Constrained genes and regions are enriched in deleterious variation, and nuclear constraint metrics such as those calculated using the Genome Aggregation Database (gnomAD) are widely used for genetic analysis of human phenotypes^1–4^. Constraint can also identify which genes and gene regions are most essential for function, given these are subject to the strongest selection^1–4^.

We recently led the expansion of the gnomAD population database to include mtDNA variation^5^. The gnomAD v3.1 dataset reports homoplasmic and heteroplasmic variants identified in 56,434 humans and at >50% of mtDNA positions^5^, representing one of the largest mitochondrial population databases. gnomAD v3.1 therefore provided an opportunity to explore mitochondrial constraint. Evidence of mtDNA selection includes pathogenic variation being rare and typically only observed at low heteroplasmy levels in the population^5^, reflecting that pathogenic variants cause disease when their heteroplasmy level is high enough to produce cellular dysfunction^7,11^. Negative selection in the human germline has also been demonstrated by reduced transmission of deleterious mtDNA variants across mother-child pairs^19–23^. While ratios of nonsynonymous to synonymous variation or variants per base have been historically used to assess mtDNA selection^24–27^, these do not account for differences in mutability between variant types and base positions, limiting their utility. Furthermore, these studies often focused on homoplasmies, observed variation, and small datasets only^19,25–27^, providing an incomplete picture. We therefore sought to assess mitochondrial constraint in gnomAD; an approach that could overcome limitations of previous studies assessing selection and provide tools to aid mtDNA analysis.

Here we describe the development of a mitochondrial constraint model, and its application to gnomAD to quantify the strength of selection across the mtDNA and within each gene. We show that mtDNA variation is subject to strong negative selection in line with its essential role in cellular function. We also demonstrate that our constraint metrics can aid the classification of mtDNA variants that contribute to human phenotypes. This work establishes the first constraint model for the human mitochondrial genome, and provides novel insights into which genes, regions and positions in the mtDNA are most essential for human health.

### Establishing a constraint model for the human mtDNA

We aimed to analyze constraint in the human mtDNA by comparing the observed level of variation in gnomAD to that expected under neutrality. The large size and diversity of the gnomAD population database, its inclusion of homoplasmic and heteroplasmic variation, depletion of severe pediatric disease, and stringent quality control makes it well-suited for the study of constraint^5^. First, we developed a mitochondrial mutational model to calculate expected variation, by using a composite likelihood model and a curated dataset of *de novo* mutations to quantify mutability in trinucleotide contexts. This likelihood model is well suited for possible sparsity of counts per context in mtDNA, and since it was developed for quantifying mutability in the nuclear exome^28^ we adapted it for analysis of mtDNA mutability (Methods). The mutational signature predicted by our model was consistent with previous reports^19,29^, showing increased likelihood of transitions over transversions, strand bias for transitions, increased likelihood of C>T when G is in the +1 position, and a distinct signature in the non-coding OriB-OriH region (Fig. 1a, Extended Data Fig. 1a). The observed level of neutral variation in each locus in gnomAD was highly correlated with their mutation likelihood (Pearson correlation coefficient R>0.99, p-value<2.2e-16), which was also a significantly better predictor of observed levels than locus length (Supplementary Methods), establishing the predictive value of the mutational model (Methods).

**Figure 1.**
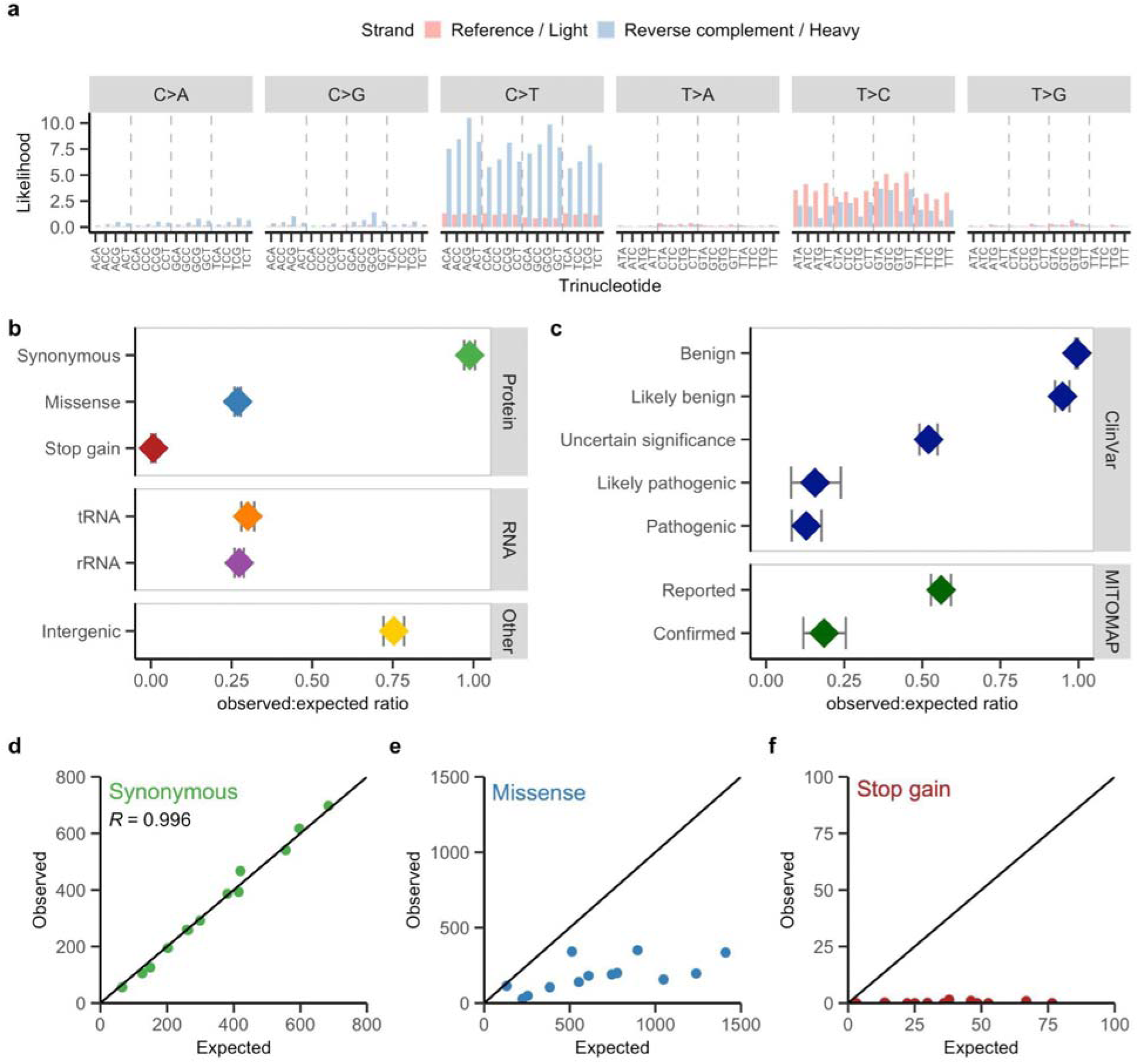
Mutability and constraint in the human mtDNA. **(a)** Trinucleotide mutational signature of mtDNA mutations predicted by the composite likelihood model. Mutation likelihoods for the six pyrimidine substitution types across 96 trinucleotides are shown, colored by whether the reference nucleotide is in the reference ‘light’ or reverse complement ‘heavy’ strand. This is computed for the reference sequence, excluding OriB-OriH (which has a distinct signature). **(b)** The observed:expected ratio of each functional class of mtDNA variation in gnomAD (synonymous, *n*=8219; missense, *n*=24,021; stop gain, *n*=1603; tRNA, *n*=4512; rRNA, *n*=7536 and non-coding, *n*=3618). Error bars represent the 90% confidence interval. **(c)** The observed:expected ratio of disease-associated mtDNA variation in ClinVar and MITOMAP databases in gnomAD, grouped by classification. 2607 ClinVar variants (benign, *n*=910; likely benign, *n*=491; uncertain significance, *n*=1000; likely pathogenic, *n*=57 and pathogenic, *n*=149) and 881 MITOMAP variants (reported, *n*=791 and confirmed, *n*=90) are included. Error bars represent the 90% confidence interval. **(d-f)** Observed and expected sum maximum heteroplasmy of variants in each protein in gnomAD are plotted for synonymous **(d)**, missense **(e)** and stop gain **(f)** variants. The Pearson correlation coefficient R is also shown in (d). Values for (d-f) are provided in Supplementary Dataset 1.

We used this mutational model to calculate the expected level of mtDNA variation in gnomAD, comparing it to the observed to quantify constraint for different functional classes of variation. Heteroplasmy is an important consideration for assessment of mitochondrial constraint given negative selection can reduce the heteroplasmy level of an observed mtDNA variant below a ‘disease threshold’. Indeed, most observed pathogenic variants in gnomAD had a maximum heteroplasmy value (expressed as a fraction, range 0.0-1.0) of <1.0, consistent with selection against high heteroplasmy^5^. We therefore calculated the observed and expected sum of maximum heteroplasmy of mtDNA variants in gnomAD, rather than the number of (unique) variants per nuclear models^1^, to capture selection against heteroplasmy (Methods). Simulation of germline mtDNA mutation and heteroplasmy drift across generations supported that mitochondrial mutation rates correlate with maximum heteroplasmy under neutrality (Extended Data Fig. 2d), in line with published work reporting that the expected number of neutral mutations drifting to homoplasmy (i.e. towards a maximum heteroplasmy of 1.0) increases linearly with mutation rate in cells^29^, corroborating the validity of using our mutational model to assess maximum heteroplasmy (Supplementary Discussion). We quantified mitochondrial constraint in gnomAD as a ratio of observed to expected variation and calculated a 90% confidence interval (CI) around these ratios (Methods). These values provide an inference on the strength of selection against a group of variants (e.g. all missense within a protein-coding gene).

Evaluation of functional classes of mtDNA variation showed that the predicted severity of each class correlated with their ratio of observed:expected variation (Fig. 1b), such that synonymous variation had a ratio close to 1 (0.99, CI 0.97-1.01) and stop gain close to 0 (0.008, CI 0.001-0.015). Other classes of variation in genes lay between these values; partial depletion of expected non-coding variation was also observed. These data show a similar selective pressure against missense, tRNA and rRNA variation, notable given the latter is typically overlooked due to the absence of *in silico* tools for predicting their effect^12^. Selection against disease-associated variation in ClinVar and MITOMAP databases, which is mostly missense and tRNA variants (Extended Data Fig. 1b-c), also correlated with classification with observed:expected ratios ranging from 0.13 for pathogenic variation (CI 0.08-0.18) to 0.995 for benign variation (CI 0.99-1.0) (Fig. 1c). Missense and tRNA variants predicted as deleterious by *in silico* algorithms were more constrained than those predicted to be tolerated, although some categories of variation predicted tolerated also appeared constrained (Extended Data Fig. 1d). Evaluation of functional classes in another large mitochondrial population database with heteroplasmy (HelixMTdb^30^) produced comparable results, supporting the robustness of our model (Extended Data Fig. 1e). Collectively, these data establish a constraint model for the human mtDNA.

### Protein gene and regional constraint identifies functionally critical sites

We evaluated gene constraint for the three major classes of protein variation; synonymous, missense, and stop gain variants. The observed and expected values for synonymous variants in each gene were highly correlated (R=0.996), consistent with their minimal selection (Fig. 1d). In contrast, stop gain variants were nearly absent in the population (Fig. 1f), as expected. This is in line with the paucity of predicted loss of function (pLoF) mtDNA variants in humans^5,30,31^, and indicates most are not compatible with life. Most proteins showed depletion of expected missense variation (Fig. 1e), and evaluation of observed:expected ratios revealed a spectrum of missense tolerance (Fig. 2a). We adopted the observed:expected ratio 90% CI upper bound fraction (OEUF) as a conservative measure of constraint akin to nuclear constraint models applied to gnomAD^1^; an approach which accounts for any uncertainty around the ratio to avoid overestimating constraint. Gene missense OEUF values ranged from 0.16 to 0.98 (Supplementary Dataset 1) and were correlated with gene function, such that complex V and (to a lesser extent) III subunit genes were most tolerant and complex I and IV least tolerant of missense (Fig. 2a). This is in line with complex I and IV defects being the most common causes of pediatric disease^32^, and with complex I, III, and IV but not complex V defects decreasing mitochondrial membrane potential, which is a trigger of mitochondrial degradation^33^. These data are also supported by studies in mtDNA ‘mutator’ mice, where analysis supported that complex V and III genes are subject to the weakest selection^24^. Comparison with conservation showed some similarities, such as *MT-ATP8* being the least conserved and least constrained, but was not significantly correlated (Extended Data Fig. 3a).

**Figure 2:**
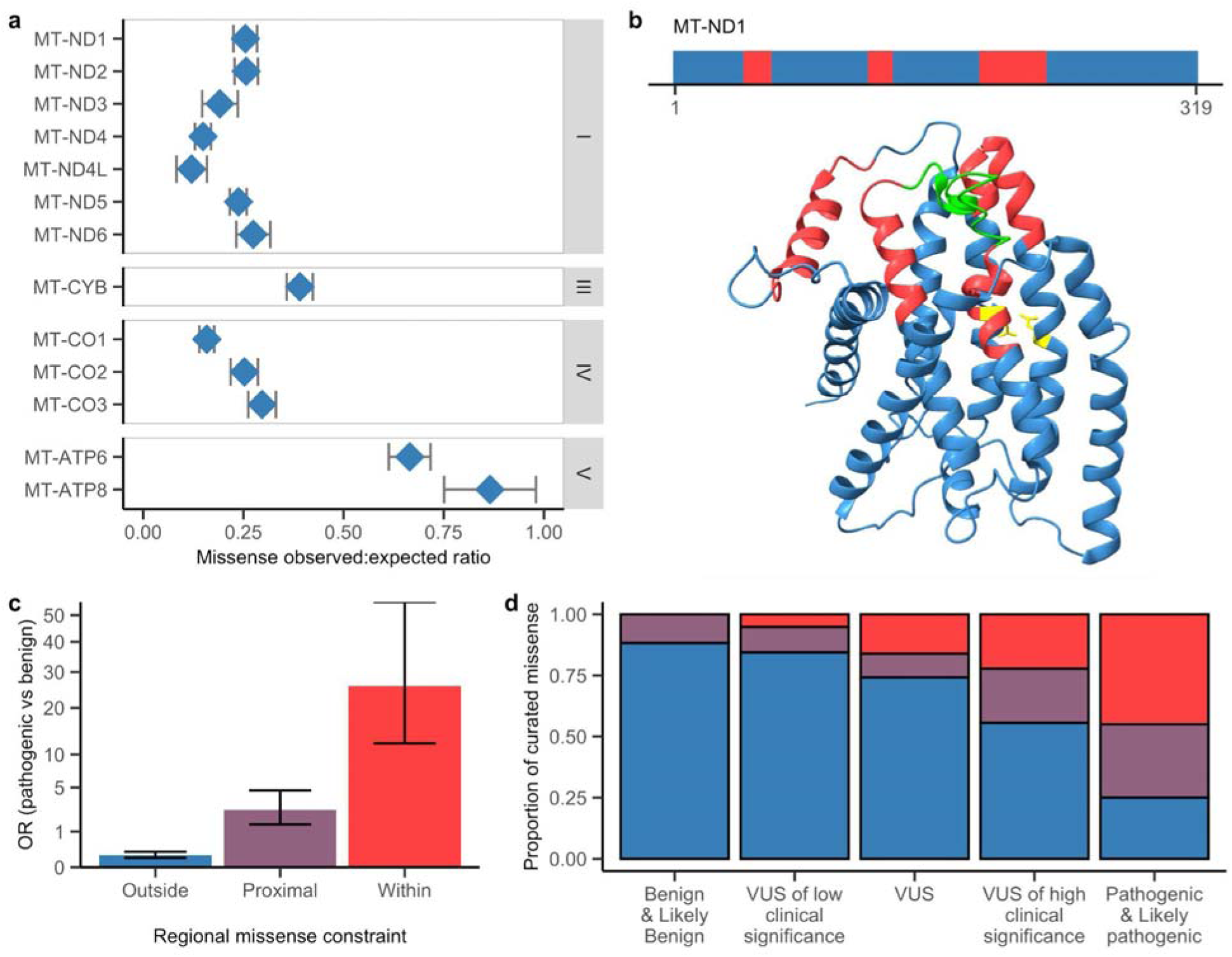
Assessment of missense constraint identifies gene and regional constraint. **(a)** The missense observed:expected ratio for each protein, ordered by the OXPHOS complex it belongs to (I, III, IV and V). Error bars represent the 90% confidence interval. Values are provided in Supplementary Dataset 1. **(b)** Areas of regional missense constraint identified in MT-ND1 are shown in red within linear protein sequence (top) and 3D protein structure (bottom). Residues in green form the shallow part of the quinone binding pocket and those in yellow are involved in proton pumping per Kampjut and Sazanov^34^. **(c)** The odds ratio (OR) enrichment of pathogenic (*n*=79) vs benign (*n*=625) missense variants (most severe consequence) outside, proximal to (<6 Ångstrom distance from), and within areas of regional constraint; the y-axis is displayed with square root scale. Error bars represent the 95% confidence interval. **(d)** The proportion of curated missense variants from a clinical genetics service that are within (red), proximal to (purple), or outside (blue) regional missense constraint, categorized by classification. 171 missense variants are included (benign & likely benign, *n*=34; VUS of low clinical significance, *n*=77; VUS, *n*=31; VUS of high clinical significance, *n*=9 and pathogenic & likely pathogenic, *n*=20). The color legend is per (c). VUS are variants of uncertain significance.

Missense tolerance can vary within proteins, a phenomenon called regional missense constraint, whereby specific gene regions can be more constrained than the gene^3^. Since these regions are enriched in pathogenic missense in the nuclear genome^3^, we developed a method to assess regional constraint in the mtDNA (Methods). We also hypothesized that regional constraint could have utility for revealing functionally critical regions, since there is a lack of functional domain annotations for these genes. Approximately 15% of total protein sequence was regionally missense constrained, and all proteins except MT-ATP8 had at least one region identified (Extended Data Fig. 4a, 5a, Supplementary Dataset 2). Mapping of regional constraint onto protein structures revealed that many were located in close proximity in 3D space, as was the case for complex I subunit MT-ND1 where regional constraint topologically clustered in the binding pocket for quinone, a molecule essential for complex I function (Fig. 2b). Manual inspection revealed other areas of regional constraint likely to be functionally critical, such as those clustering at the heme and quinone sites in MT-CYB (Extended Data Fig. 5d-e, Supplementary Discussion). Although functional domain annotations aren’t readily available, there are residues of known functional importance (i.e. cofactor binding or proton transfer). These residues were highly missense constrained, having an OEUF value (0.06) comparable to that for pLoF variants, illustrating their critical role in function. Approximately 60% of these functional residues were located in areas of regional constraint or were proximal to it within 3D space (Extended Data Fig. 5a), indicating these intervals are enriched in functionally critical sites. These data also highlight the utility of regional constraint to identify residues not realized as functionally important.

To determine if there was an association between regional constraint and pathogenic variation, we calculated the odds ratio (OR) using pathogenic missense variants reported in ClinVar or MITOMAP versus benign ClinVar variants. Regional constraint was highly enriched in pathogenic versus benign missense (OR=26, 95% CI=12-55); residues in close proximity to regional constraint in 3D space (<6 Ångstrom distance) were also enriched in pathogenic to a lower extent (OR=2.6, 95% CI=1.4-4.6) (Fig. 2c). This was supported by mtDNA variants curated by a clinical genetics service in cases suspected to have mitochondrial disease, where the proportion of missense variants within or proximal to regional constraint correlated with classification (Fig. 2d). Regional missense constraint performed particularly well at discriminating ‘true negatives’, given the lack of benign variants within regional constraint (Fig. 2d, Extended Data Fig. 5b). Regional constraint therefore provides a tool for variant classification, including for utilization of the ACMG pathogenic criterion PM1 “*Located in a mutational hotspot and/or critical and well-established functional domain without benign variation*” which was omitted from mitochondrial ACMG/AMP variant classification guidelines due to lack of applicability^12^. Overall, we demonstrate the utility of mitochondrial regional constraint for variant classification, and establish it as a tool for prioritizing variants of uncertain significance in individuals with mitochondrial diseases.

### Revealing constraint across and within mitochondrial RNA genes

Unlike the nuclear genome, most genes in the mtDNA encode for RNAs; specifically tRNAs and rRNAs required for mitochondrial translation. The tRNAs showed a spectrum of intolerance to base substitutions, with OEUF values ranging from 0.21 for *MT-TM* to 0.87 for *MT-TT* (Fig. 3a, Supplementary Dataset 1). These gene constraint values were not significantly correlated with each tRNA’s codon usage in the mtDNA (Extended Data Fig. 3c), suggesting other factors are driving selection. Indeed, *MT-TM* is the most constrained tRNA, encoding the initiator (and elongator) tRNA^Met^, while the tRNA with the second lowest OEUF (*MT-TL1*) is also notable for encoding the binding sequence for the mitochondrial transcriptional terminator MTERF1^35^. Gene conservation scores only partially correlated with constraint (Extended Data Fig. 3b). The low OEUF values observed for the two rRNA genes (OEUF 0.28 and 0.30) are striking given these loci are typically overlooked in analyses, due to a lack of *in silico* tools for predicting rRNA variant effect^12^. However our data is consistent with prior studies reporting reduced transmission or frequency of rRNA variation^5,19,20,25^, supporting these genes likely harbor undiscovered pathogenic variation.

**Figure 3:**
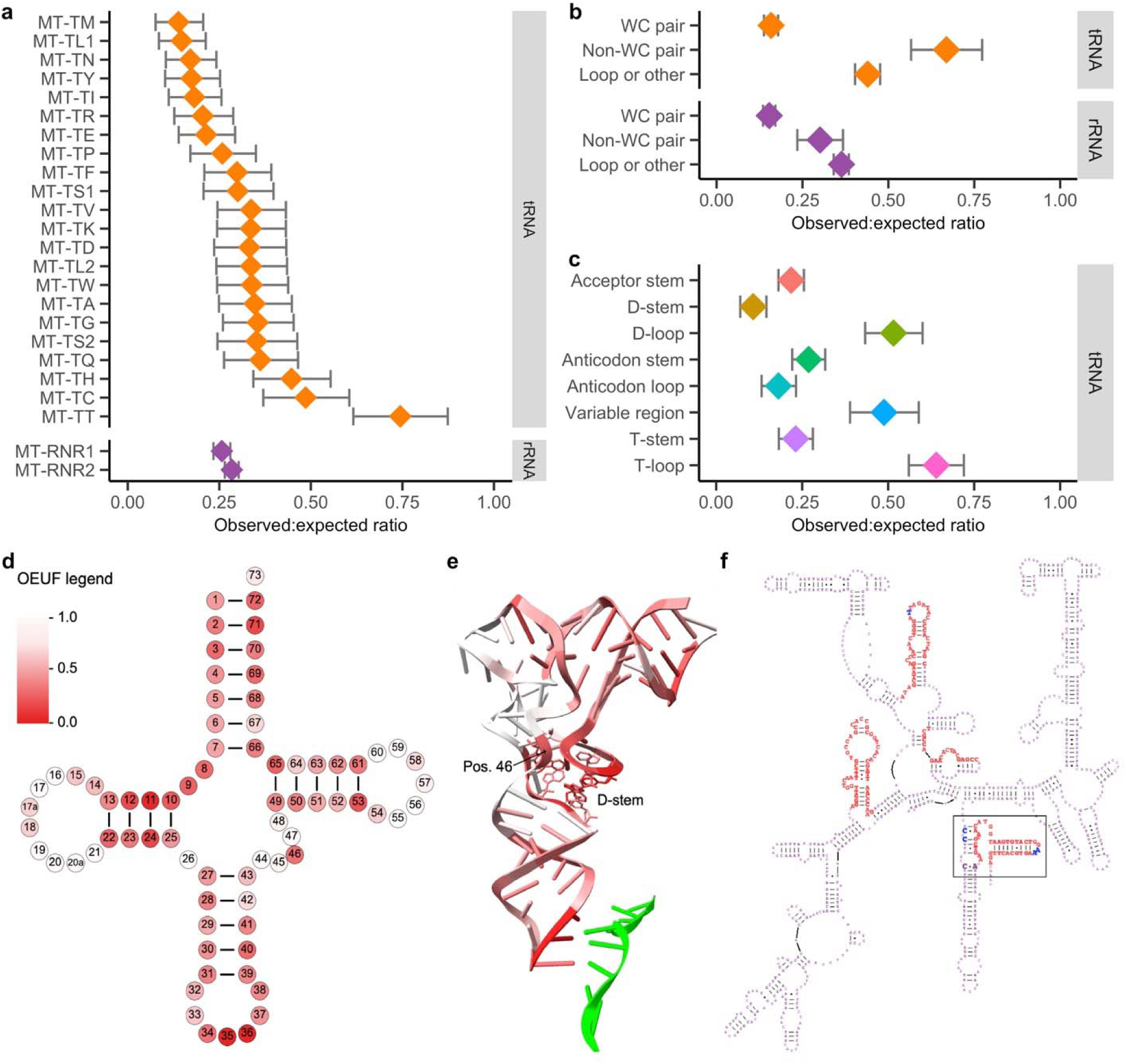
Constraint across and within RNA genes. **(a)** The observed:expected ratio for variants in each RNA, ordered by RNA type and value. Values are provided in Supplementary Dataset 1. **(b)** The observed:expected ratio for each base type in tRNA (WC pair, *n*=2364; non-WC pair, *n*=318 and loop or other, *n*=1842) and rRNA (WC pair, *n*=3078; non-WC pair, *n*=354 and loop or other, *n*=4104). WC represents Watson-Crick, and loop or other includes all single-stranded regions. **(c)** The observed:expected ratio for each tRNA domain (acceptor stem, *n*=933; D-stem, *n*=468; D-loop, *n*=411; anticodon stem, *n*=672; anticodon loop, *n*=462; variable region, *n*=270; T-stem, *n*=606 and T-loop *n*=459); their secondary structure location is per Extended Data Fig. 6c. Error bars in (a-c) represent the 90% confidence interval (CI). **(d)** The observed:expected ratio CI upper bound fraction (OEUF) of variants at each tRNA secondary structure position. Darker colors represent lower values, per the legend. Values are provided in Supplementary Dataset 4. **(e)** Each tRNA position OEUF mapped onto tRNA tertiary structure; color legend is per (d). Labeled position 46 and D-stem positions are shown in nucleotide style. The mRNA molecule is colored green. **(f)** Areas of regional constraint within MT-RNR1 secondary structure, indicated by red font. The box highlights an area including regional constraint, modified bases (blue font) and disease-associated variants (at m.1494 and m.1555, bold purple font); also shown in tertiary structure in Extended Data Fig. 5g.

The RNAs form secondary structures with double-stranded stems and single-stranded loops. Variants disrupting Watson-Crick (WC) base pairs within stems were highly constrained, at similar levels in tRNAs and rRNAs (OEUF 0.18 and 0.17, Fig. 3b). This is in line with nearly 70% of pathogenic tRNA variants breaking WC pairs (Extended Data Fig. 6a), and supports that this variant type may especially harbor deleterious variants in rRNA genes. Non-WC pairs in stems and single-stranded loops were more tolerant of variation than WC pairs, especially within tRNAs (Fig. 3b). Post-transcriptionally modified bases were also more constrained than non-modified bases, consistent with their role in RNA stability and function^36^ (Extended Data Fig. 6b). The tRNAs share a cloverleaf secondary structure, which has annotated domains (Extended Data Fig. 6c). Assessment across domains revealed clear differences in tolerance to variation, such that the D-stem was most constrained (OEUF 0.15) and the T-loop the least (OEUF 0.72, Fig. 3c). These values correlated with domain enrichment in pathogenic variants (Extended Data Fig. 6d), and are consistent with studies noting increased pathogenic burden in some of the most constrained domains, especially the anticodon and acceptor^25,37^. Although the latter have obvious functional significance due to binding mRNA or amino acids, these data also highlight a critical role for the D-stem which is involved in the formation of the L-shaped tertiary tRNA structure^38^.

Owing to the shared structure of the tRNAs, each base can be assigned a position number. Evaluation across each tRNA position revealed a spectrum of constraint (Fig. 3d, Supplementary Dataset 4). For example, the non-wobble anticodon positions (35-36) and position 11 in the D-stem were highly intolerant of variation, with values similar to pLoF variants (OEUF <0.05), while position 26 between stems was tolerant of variation (observed:expected ratio 1.04 with OEUF 1.35). These data affirm the pLoF effect of non-wobble anticodon variants^29,39^, whilst also highlighting positions not widely appreciated as functionally important. This includes the position 11-24 pairing in the D-stem, as well as position 46 in the variable region that interacts with the D-stem in the tertiary structure (Fig. 3e). Unlike the tRNAs, the rRNAs do not share a common structure. Therefore, we assessed regional constraint to characterize tolerance to variation across each rRNA (Supplementary Dataset 2). Approximately 15% of rRNA bases were regionally constrained (Fig. 3f, Extended Data Fig. 4b, 5c, 5f). Post-transcriptional modifications and rRNA:rRNA bridges connecting the mitoribosomal subunits have a critical role in rRNA function^36,40^. Accordingly, 70% of modified or intersubunit bridge bases were within regional constraint or in close proximity to it in the tertiary rRNA structure (Extended Data Fig. 5c), indicating these intervals are enriched in functionally critical sites. Indeed, the most constrained rRNA region encodes a site involved in tRNA binding during translation with an OEUF comparable to protein gene pLoF OEUF values (Supplementary Discussion). The two well-established rRNA pathogenic variants, that cause deafness^41^, were also nearby to regional constraint (∼10 Ångstrom) (Fig. 3f, Extended Data Fig. 5g). Collectively, these data identify which RNA sites are most important for function, and thus most likely to harbor deleterious variation.

### Non-coding elements involved in mtDNA replication and transcription are constrained

Approximately 10% of the mtDNA is non-coding. Most of this sequence lies within the ‘control’ region, which contains annotated elements involved in mtDNA transcription and translation^42^. Since we observed partial depletion of expected non-coding variation (Fig. 1b), we calculated constraint metrics for non-coding elements (Supplementary Dataset 5). Several elements in the control region were constrained (Fig. 4); this included the promoter for transcription of the light strand (LSP, OEUF 0.53) and conserved sequence block 3 (CSB3, OEUF 0.33). CSB3 was recently shown to be essential for the function of 7S RNA, a key regulator of transcription initiation in the mitochondria^43^. In contrast, the hypervariable sequences (HVS1-3) within the control region showed OEUF values ranging from 0.90-1.09; these regions are known to be highly polymorphic, and therefore were expected to be tolerant of variation. The hypervariable sequence with the lowest OEUF value (HVS2) was also the one that encoded the most annotated elements (Fig. 4).

**Figure 4:**
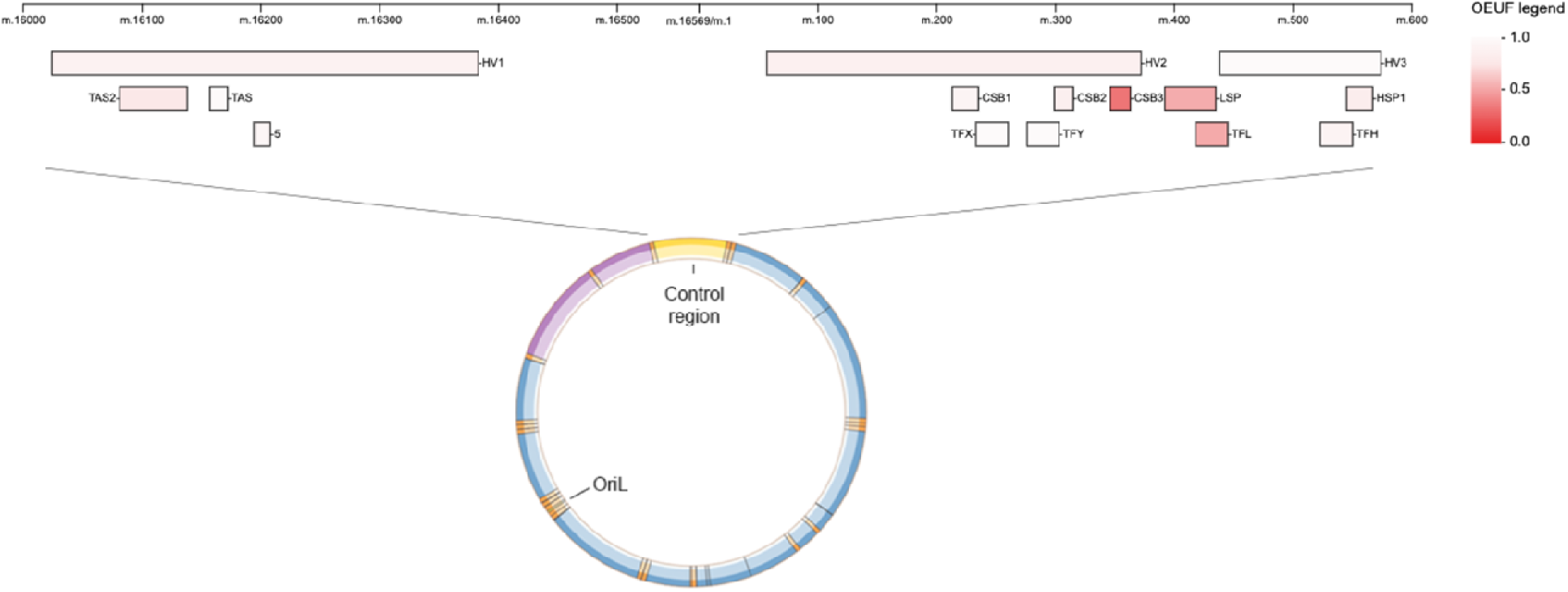
Measuring constraint across non-coding elements. Top schematic shows annotated elements within the non-coding control region, which spans the artificial chromosome break (m.16569-1). The top row includes the three hypervariable sequences (HV1, HV2, HV3), the second row includes termination-associated sequences (TAS, TAS2), conserved sequence blocks (CSB1, CSB2, CSB3) and L-strand and H-strand promoters (LSP, HSP1), and the third row includes a control element (MT-5) and transcription factor binding sites (TFX, TFY, TFL, TFH). The observed:expected ratio 90% confidence interval upper fraction (OEUF) within each element is shown per the color gradient legend; darker colors represent lower OEUF. Values are provided in Supplementary Dataset 5. The bottom schematic shows the position of the control region and origin for replication of the light strand (OriL) within the mtDNA, with encoded loci colored by their type (non-coding in yellow, protein blue, rRNA purple and tRNA orange).

The non-coding sequence outside of the control region encodes an element of functional significance, the origin for replication of the light strand (Fig. 4), which was also constrained (OriL, OEUF 0.38). The OriL overlaps two tRNA genes (*MT-TN* and *MT-TC*) but the non-coding section encoding the initiation site was also markedly depleted of expected variation (OEUF 0.30), consistent with its essential role in mtDNA maintenance^44^. These data support that some non-coding variants are likely deleterious due to effects on mtDNA replication and transcription.

### Characterizing the most constrained sites in the human mtDNA

To identify the most constrained sites within the entire mtDNA, agnostic of locus annotation, we assessed local intolerance to base or amino acid substitution (i.e. missense only in protein genes) at and around every position using an overlapping sliding window method (Methods). This provided a mean OEUF value for every position, where bases in the closest proximity contribute more of the signal, which was percentile ranked to derive a score between 0-1 termed the mitochondrial local constraint (MLC) score. The most locally constrained positions with a score >0.99 include residues in MT-CO1 and MT-CO2 binding copper, notable given copper metalation is required for complex IV function^45^, as well as residues in the MT-ND4 binding site for complex I inhibitor rotenone^34^ (Fig. 5a). Numerous RNA bases had high scores >0.95, including rRNA sites involved in tRNA and mRNA binding during translation^46^ (Fig. 5a, Supplementary Video). Non-coding bases were depleted from the highest score quartile (0.75-1.00) (Fig. 5b), and those with the highest scores were in the origin for light strand replication or focally distributed in the control region (Extended Data Fig. 7a). The latter included a recently discovered light strand promoter^47^ and two regions of unknown function, including one between m.16,400-16,450 previously reported to be depleted of heteroplasmic variation^19^, and another across the artificial chromosome break. These two regions also had a lower population frequency of variants (Extended Data Fig. 7b-d), although more work is needed to determine what is driving their signals (Supplementary Discussion).

**Figure 5.**
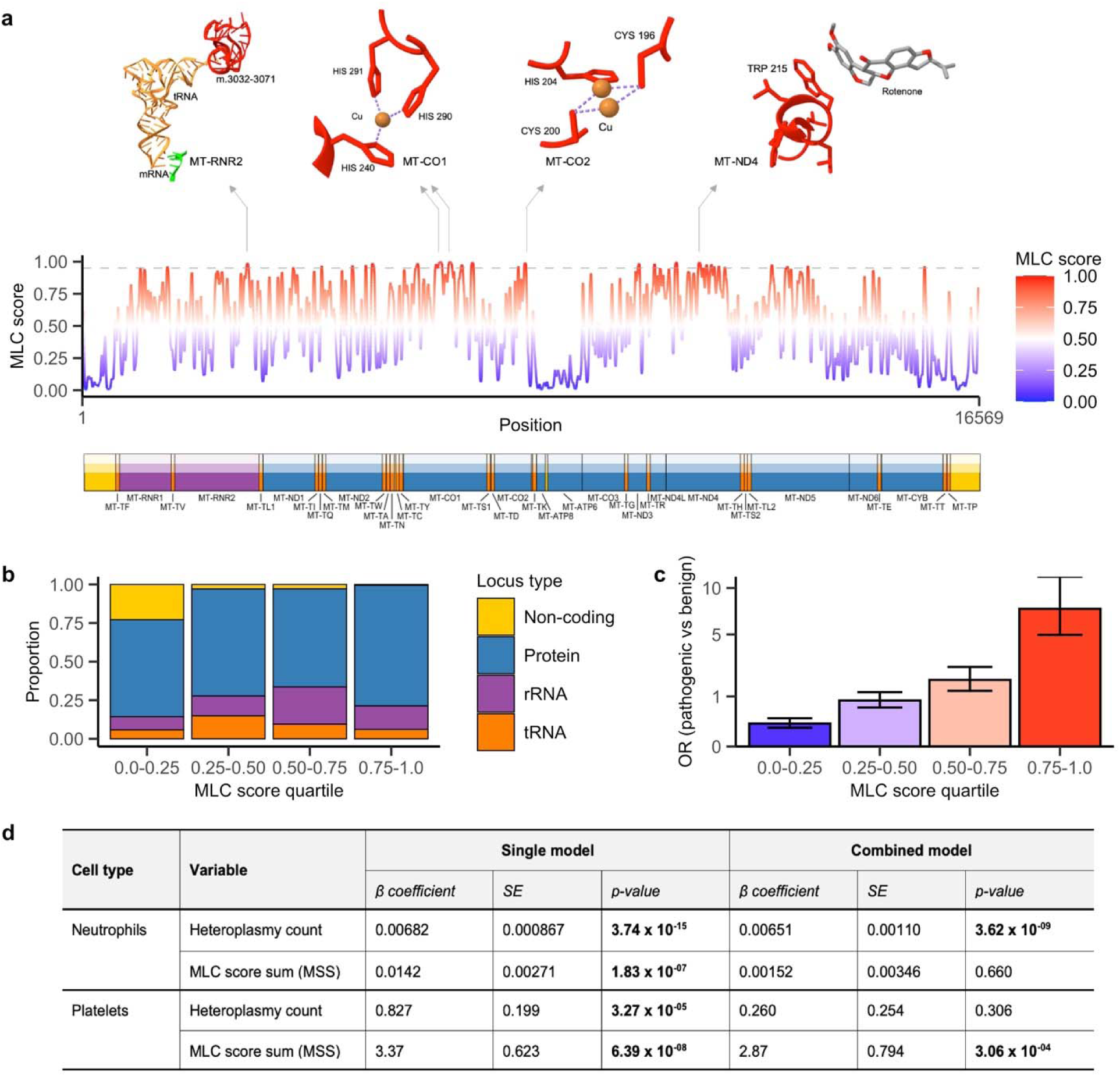
Assessment of mitochondrial local constraint (MLC) scores. **(a)** The MLC score across every base position in the human mtDNA. Encoded genes are shown below, colored as follows: protein in blue, rRNA purple, tRNA orange, and non-coding yellow. The dashed gray line represents a score of 0.95. Per base scores are provided in Supplementary Dataset 6. The top panel shows examples of positions with the highest scores, from left to right in MT-RNR2 at the tRNA/mRNA interface in the mitoribosome (m.3032-3071), copper binding sites in MT-CO1 (p.240, p.290-291) and MT-CO2 (p.196, p.200, p.204), and residues in the MT-ND4 rotenone binding site (e.g. p.215). **(b)** The proportion of bases in each locus type in each MLC score quartile. **(c)** The odds ratio (OR) enrichment of pathogenic (*n*=205) vs benign (*n*=884) variants within each MLC score quartile, for RNA base and amino acid substitutions only. Error bars represent the 95% confidence interval. **(d)** Table showing β coefficients, standard error (SE), and p-values from linear regression models of the association between platelet or neutrophil count and heteroplasmy count and or MLC score sum. Regressions were run as separate models (‘Single’) or together in the same model (‘Combined’), and adjusted for age, sex and smoking status. Significant p-values are bolded.

Variants at positions with high MLC scores are predicted to be more deleterious. To investigate this, we assessed the odds ratio (OR) enrichment of pathogenic vs benign variants reported in ClinVar and MITOMAP databases across score quartiles, for RNA and missense variants. Pathogenic variants were 7.5 times more likely to be within the highest score quartile than benign variants (95% CI=4.96-11.5, Fig. 5c); they were also enriched across scores between 0.50-0.75 (OR=1.8, 95% CI=1.2-2.5) and depleted from the lowest quartile (OR=0.21, 95% CI=0.14-0.32). Although depleted, some confirmed pathogenic variants were in the lowest score quartile (Extended Data Fig. 8a-d). Since this score measures local constraint around each position, pathogenic variants can have a low score if their neighboring positions are tolerant of variation; conversely benign variants can have a high score if their neighbors are intolerant of variation. Therefore, while a higher score increases the likelihood of pathogenicity, it does not preclude benign impact (Supplementary Discussion). Base positions with the highest scores were highly intolerant of indel variants, supporting that variation at these sites is more likely to impair mitochondrial function (Extended Data Fig. 8e); they were also more likely to be conserved across vertebrates (Extended Data Fig. 8f). Since the score measures missense tolerance specifically in protein genes, we also assigned non-missense protein SNVs scores to extend application to all SNVs (Methods). SNVs with higher scores were more likely to be seen only as heteroplasmies and ultra-rare homoplasmies in several mtDNA population databases^5,30,31^ (Extended Data Fig. 8g-i), in line with their increased pathogenicity^5,12^. Overall, these data support that MLC scores can serve as a predictor of deleterious impact, across every locus type in the mtDNA.

We were interested to use the MLC score to assess the impact of mtDNA variation on phenotypes in ∼200,000 individuals with genome sequencing data in the UK Biobank. Previous studies had demonstrated an association between mtDNA copy number and blood cell counts^48^, and we likewise observed an association between heteroplasmic variant counts and blood cell counts in study participants, specifically for neutrophils and platelets (p-values 3.74×10^-15^ and 3.27×10^-05^), using a linear regression model adjusted for age, sex, and smoking status (Fig. 5d). We generated a MLC score sum (MSS) for each participant from all of their heteroplasmies, to assess their functional impact. When both heteroplasmy count and MSS were included in the same model, neutrophil count was only significantly associated with heteroplasmy count (p-value 3.62×10^-09^), while platelet count was only significantly associated with MSS (p-value 3.06×10^-04^) (Fig. 5d). These results support different roles for mitochondria in neutrophils and platelets, with mtDNA likely playing a causal role in platelet count and non-causal role in neutrophil count, consistent with the lack of nuclear DNA in platelets and the diminished role of mitochondria in neutrophils (Supplementary Discussion). Further analysis revealed significant associations between MSS and other phenotypes, including mortality, as described in an accompanying manuscript^49^. These data support the utility of the MLC score, and provide examples of how these metrics can provide novel insight into the role of mtDNA variation in human phenotypes.

## Discussion

Mitochondrial DNA is an essential, yet often overlooked, part of the human genome. In this manuscript, we advance efforts to map constraint across the human genome by expanding constraint models to include the mtDNA. Application of our model to gnomAD revealed strong depletion of expected variation, supporting that only a fraction of total deleterious variation in the mtDNA has been characterized. Our constraint metrics provide tools to help address this gap in knowledge, by identifying which genes, gene regions, and positions across the human mtDNA are most likely to harbor undiscovered pathogenic variation. We validated the utility of these metrics by using disease-associated variants and functional annotations, and provided examples of how they can be used to investigate the role of mtDNA variation in rare and common phenotypes. Given the mtDNA is emerging as an important contributor to a myriad of phenotypes, we anticipate that these data will be a widely useful resource for studies that aim to elucidate the contribution of mtDNA variation to disease and other health outcomes.

Strikingly, we identified constraint in genes and regions commonly overlooked in disease analyses. This includes the rRNA genes, which are critical for translation but often not evaluated due to a lack of *in silico* tools, as well as non-coding elements involved in mtDNA replication or transcription where variation is typically assumed benign. Although surprising, these data are supported by studies reporting reduced transmission or frequency of variants in these loci^5,19,20,23,25^, indicating they likely harbor undiscovered pathogenic variation. Our exploration of constraint within the mitochondrial rRNA and tRNA genes may further offer a framework for the assessment of their nuclear-encoded counterparts given constraint within RNA genes has been little explored, though recent work establishing a mutational model for these genes will pave the way for this^50^. We also identified strong depletion of expected variation in protein genes with few pathogenic variants reported, such as *MT-CO1* which harbored some of the most missense constrained sites in the mtDNA and yet has no ‘confirmed’ pathogenic missense^31^. This could reflect a bias towards the study of genes that already have well-established pathogenic variants, since genes with the most ‘confirmed’ pathogenic variants are enriched with those most frequent in the population^5,31^, and or an increased burden of deleterious variants not compatible with life.

Our work capitalizes on the availability of heteroplasmy data within gnomAD; an inclusion that represents a significant advance for the study of mtDNA population genetics. Importantly, this enabled us to capture selection against both variant occurrence and heteroplasmy in gnomAD, the latter representing a phenomenon unique to the mtDNA. However, there are some important considerations with using gnomAD to assess mitochondrial constraint. Most gnomAD samples are from blood^5^, a dividing cell type that can exhibit lower heteroplasmy of pathogenic variants than other (post-mitotic) tissues due to selection across cell divisions^51^. The depletion of expected variation we observed may therefore be greater than in other tissues, although selection in the germline and embryos is stronger than in somatic cells^22,23^. Future studies assessing constraint in tissue datasets, as well as for inherited vs somatic mutations, will shed light on tissue-specific selective forces shaping mtDNA variation. Furthermore, the European overrepresentation in gnomAD may bias our metrics to have the highest predictive value for European haplogroups. This is relevant given reports of variants only causing mitochondrial dysfunction in specific mtDNA haplogroup backgrounds^12,52^, and of nuclear genetic background impacting heteroplasmy transmission^19^. Ongoing expansion of gnomAD can address this, as well as provide the power to analyze haplogroup and population differences in constraint.

It is important to note that our constraint model is specifically capturing negative selection against variants that have functional impacts at heteroplasmy; a criterion the majority of reported pathogenic variants meet^31^. Therefore these metrics are best suited for assessment of heteroplasmic variation. Future iterations of our model which address challenges with incorporating population frequency data will be needed to improve utility for detecting weaker selection against homoplasmic variation. Despite this caveat, our results support that mtDNA selection is stronger than previously appreciated. This likely reflects limitations of methods previously used to assess selection in human population datasets, including assumptions that the expected number of variants across genes should equal the observed, incomplete separation of missense and synonymous variants, and assumption of equal mutability per base pair^20,21,24–26^. We observed a spectrum of intolerance to variation across the mitochondrial genome, establishing a map of the regions that are likely to be most critical for mitochondrial function.

Given the lack of functional domain annotations for most of the coding and non-coding sequence in the human mtDNA, our data provides a rich set of candidate sites whose characterization will provide novel insights into mitochondrial genome function.

Selection against mtDNA variants is complex, representing a combination of forces at an organismal (i.e. reduced survival and reproductive fitness), cellular (i.e. germline or somatic), and even organelle level (i.e. mitochondrial)^7^. While additional research is needed to tease apart the contribution of each to mitochondrial constraint in the human population, the near absence of expected pLoF variation in gnomAD supports that mtDNA variants with the most severe impact are not compatible with life and thus removed by selection in the germline or *in utero*. This also demonstrates the high efficiency of mechanisms for preventing accumulation of deleterious mtDNA variants in the population, necessary given its high mutation rate^22,23^. Accordingly, most pathogenic variants characterized to date will have only mild to moderate effects, given they are compatible with life but not health^53^; a point which has implications for *in silico* predictors trained on reported pathogenic variants. Studies which clarify the impact of variants on germ cell and *in utero* development, such as those occurring at the most constrained sites, will be needed to characterize both the observed and unobserved deleterious variation in the human population.

The increasing affordability of genome sequencing, coupled with the development of other mtDNA tools such as base editors^54^, are poised to usher in an exciting era of mtDNA research. In this spirit, our mitochondrial constraint model provides a tool to complement these advances and empower research into the role of mtDNA variation in human health and disease.

## Figure legends

Each figure legend is presented underneath its figure in the main text for initial submission.

## Methods

### Mitochondrial mutational model

*Mitochondrial composite likelihood model:* We adapted a composite likelihood model described by Dietlein *et al*^28^, and applied it to *de novo* mutations to quantify mutability in trinucleotide contexts in the human mtDNA. The composite model decomposes the mutational likelihood of each trinucleotide context into multiplicative factors, namely the effects of the reference nucleotide, mutation class, and flanking nucleotides, making it optimal for the possible sparsity of counts per context in the smaller mtDNA. We adapted the model for its application to the mtDNA; a detailed description is in the Supplementary Information, and summarized here.

We classified 12 base substitution types *t* ∈ {*C* > *A*, *C* > *G*, *C* > *T*, *T* > *A*, *T* > *C*, *T* > *G*, *G* > *T*, *G* > *C*, *G* > *A*, *A* > *T*, *A* > *G*, *A* > *C*} and their reference nucleotides *n*(*t*) ∈ {*A*, *C*, *G*, *T*}, and categorized them into three mutational classes *c*(*t*) ∈ {I, II, III}: transversions type I (class I), transversions type II (class II) and transitions (class III). We counted the *de novo* mutations of each base substitution type *t* to produce vector *v^type^*. The likelihood ratio of each reference nucleotide *λ_n_* where *n* ∈ {*A*, *C*, *G*, *T*} was calculated as

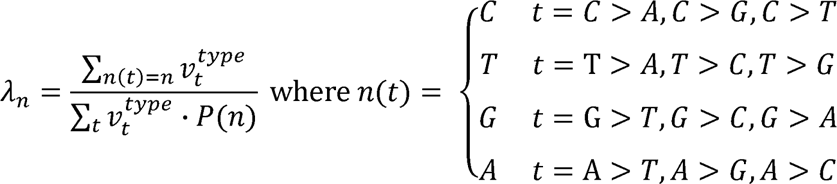

where the probability of the reference nucleotide is its frequency in the reference sequence. The likelihood ratio of each mutation class *λ_c_* was calculated as

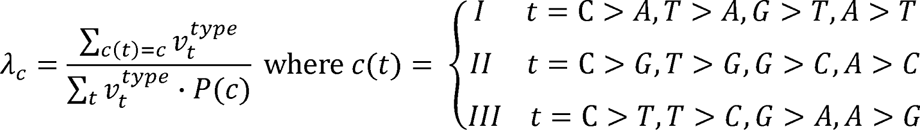

where the probability of the mutation class was derived from the relative frequency of nucleotides in the reference sequence. The likelihood ratio of each base substitution type *t* was calculated as

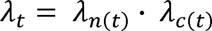

and the likelihood ratios for sequence context *λ_t,p,n_* as

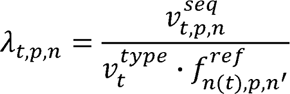

For *c*(*t*) = III, 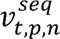 is the count of nucleotide *n* at position *p* ∈ [-1: 1]\{0} around base substitution of type *t*, and 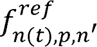 is the frequency of nucleotide *n*’ at position *p* ∈ [-1: 1]\{0} around the reference nucleotide *n*(*t*) in the reference sequence. While there is replicative strand bias for transitions (class III), this has not been established for transversions^29,55^. Therefore for *c*(*t*) = I, II the base substitution types *t* were first classified into their pyrimidine type *pyr*(*t*) due to their lower counts, and 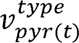 was used to count *pyr*(*t*), 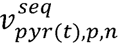 to count of nucleotide *n* at position *p* ∈ [-1: 1]\{0} around ‘*pyr*(*t*), and 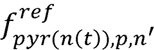 to represent the frequency of nucleotide *n*’ at position *p* ∈ [-1: 1]\{0} around the pyrimidine reference nucleotide. We then computed the mutability of each mutation class *c* at every base in the reference sequence, excluding the non-coding OriB-OriH region, for *m* ∈ [192:16196] with reference nucleotide *n*_0_ and flanking nucleotide *n_p_* at position *p* as

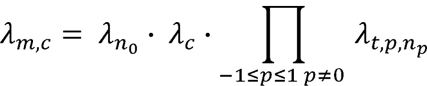

This produced a composite mutation likelihood for every possible base substitution in the mtDNA. Mutability in the non-coding OriB-OriH region *m* ∈ [1-191, 16196-16569] was quantified separately to handle its inverted signature for transitions^29^ (Supplementary Methods).

*De novo dataset:* We quantified mutability using mitochondrial *de novo* mutations from the literature and an in-house dataset. Published *de novo* mutation datasets were identified from a search of the literature^19–22,56,57^; these included germline as well as somatic *de novo* mutations which have a highly similar mutational signature in the mtDNA^29,56^. The in-house dataset of *de novo* mutations was identified from 1690 mother-child pairs with unaffected status in the SPARK cohort^58^, using genome sequencing data obtained from SFARI Base (application #12267.2). The GATK Mutect2 pipeline was used to call variants in the SPARK pairs, and variants at >1% heteroplasmy level that were present in the child but not in the mother after stringent filtering were regarded as *de novo*, akin to criteria used by others^19^. Checks were performed to confirm that *de novo* mutations across sources had highly similar mutation characteristics, and one outlier group was excluded (see Supplementary Methods). A final dataset of 4216 *de novo* mutation counts was used, extracted from each source using custom scripts run in Python v3.10. Additional details on the *de novos* and their curation are in the Supplementary Information.

*Validation:* The predictive value of the mutational model was validated by measuring the correlation between the mutation likelihoods and observed level of neutral variation in gnomAD. Haplogroup variants supplemented with variants at non-conserved sites in the lowest decile of PhyloP were used to represent neutral variation. Custom scripts implemented in Python v3.10 were used to sum the observed maximum heteroplasmy of neutral variants in gnomAD and their mutation likelihood scores across each locus. Linear regression models fitting mutation likelihoods and observed neutral variation, and their Pearson correlation coefficients and p-values, were calculated using R v3.6.1. The highly mutable G>A and T>C variants were fit separately, akin to how CpG transitions are handled separately in nuclear models^1^, as was the non-coding OriB-OriH region. Additional details on this method are in the Supplementary Information.

### Assessment of mitochondrial constraint

We measured mitochondrial constraint in gnomAD as a ratio of observed to expected variation. Specifically, we calculated the observed and expected sum maximum heteroplasmy of variation, to capture selection that occurs against the number and heteroplasmy level in the mitochondria. The observed value for a variant class and or locus in gnomAD v3.1^5^ was determined by summing the maximum heteroplasmy value of every possible SNV in the group (e.g. all missense within a protein gene). The expected value under neutrality was determined by summing the mutation likelihoods of every possible SNV in the group, and applying the linear models fit on the mutation likelihoods and observed neutral variation in gnomAD (described above). We also calculated the 90% confidence interval (CI) around each observed:expected ratio using a beta distribution, adapting the method used for nuclear constraint models^1^. Custom scripts run in Python v3.10 were used to calculate ratios and their CIs. The observed:expected ratio 90% CI upper bound fraction (OEUF) was used as a conservative measure of constraint. Additional details on these methods are in the Supplementary Information.

### Simulation of germline mtDNA mutation and heteroplasmy

To validate a correlation between mitochondrial mutation rates and population maximum heteroplasmy, we adapted a computational model by Colnaghi *et al* to simulate mutation and heteroplasmy drift for neutral mutations in the human female germline^59^. Heteroplasmy levels were tracked across 10,000 maternal lineages for five generations for 10 mutation rates (between 10^−9^-10^−7^ per base pair), using model parameters drawn from human data by Colnaghi *et al*^59^. The maximum heteroplasmy distribution for each mutation rate in the simulated population was evaluated through bootstrapping. This simulation was implemented using a custom R script (run in v.3.6.1); a detailed description of this method is in the Supplementary Information.

### Gene, non-coding, and variant *in silico* annotations

A ‘synthetic’ VCF with every possible SNV in the human mtDNA reference sequence NC_012920.1, and their Ensembl Variant Effect Predictor gene and consequence annotations, generated as described^5^ was used for computing observed and expected variation. Note that each human mtDNA gene has only one transcript, thus distinction between canonical and non-canonical transcripts was not required. Non-coding elements in the human mtDNA and their coordinates were downloaded from MITOMAP^31^; elements with expected value <10 were excluded from analyses. The coordinates for the non-coding ‘OriB-OriH’ region are per Ju *et al*^29^. Haplogroup variants were extracted from PhyloTree Build 17 as previously described^60^. The phyloP conservation scores derived from 100 vertebrate genomes, and APOGEE, HmtVar and MitoTip *in silico* predictions, were retrieved as described^5,60^.

### Protein and RNA annotations

Functional sites within the proteins were gleaned from UniProt and the literature as follows. UniProt Knowledgebase annotations were downloaded from the UniProt FTP site (https://www.uniprot.org/downloads), and binding site annotations for human mtDNA-encoded proteins were extracted from available bed files (date of access November-16-2020)^61^. Residues involved in complex I proton transfer were curated based on ovine data reported by Kampjut and Sazanov^34^; the reported residue positions were manually confirmed as equivalent to human except for MT-ND6 which was shifted by one residue. For RNA genes, the base type annotation (e.g. base pair in stem) was determined using a custom script in Python v3.10 and manually curated secondary structure data reported previously^60^. Modified bases in RNA genes and tRNA domain annotations were obtained as described^60^. The tRNA secvvvvvvgondary structure position numbers and their corresponding mtDNA position were obtained from Sonney *et al*^62^. Codon usage of each tRNA was determined by counting the corresponding amino acid within the protein-coding sequence using custom scripts in Python v3.10; for Leucine and Serine the codon sequence was used to distinguish between the two tRNAs for these amino acids. Bases involved in rRNA:rRNA bridges connecting the two mitoribosomal subunits are per Amunts *et al*^40^.

### Population databases

Data from the Genome Aggregation Database (gnomAD) v3.1 generated from whole genome sequences from 56,434 individuals were retrieved from the gnomAD browser (https://gnomad.broadinstitute.org/downloads)^5^. HelixMTdb data generated by proprietary exome+ assay of 195,983 individuals were downloaded from the Helix website (https://www.helix.com, version dated 03-27-2020)^30^; observed maximum heteroplasmy of homoplasmic variants was not reported and therefore assigned as 1.0. MITOMAP data generated from 56,910 GenBank sequences were downloaded from the MITOMAP website (https://www.mitomap.org/MITOMAP/resources, polymorphism table, download date 07-14-2022)^31^. Note that the MITOMAP database does not include heteroplasmy information.

### Disease-associated variation

Disease-associated mtDNA variants were obtained from ClinVar and MITOMAP databases. For ClinVar, all mtDNA SNVs were retrieved (download date 05-25-2022)^63^. Variants listed only to be associated with cancer were excluded to focus on germline conditions, and those with conflicting interpretations were also excluded. For MITOMAP, all disease-associated variants were retrieved (disease table, download date 05-25-2022)^31^. A total of 2607 ClinVar variants and 882 MITOMAP variants were used. For Extended Data Fig. 8d, all variants with a confirmed status and plasmy status of ‘-/+’ in MITOMAP (associated with disease at heteroplasmy only) were reviewed. Any variants reported to be observed at homoplasmy in an individual in at least one publication were shown in the ‘at homoplasmy’ group in Extended Data Fig. 8d, as per Supplementary Dataset 8. Curated missense variants identified in cases suspected to have a mitochondrial disease (in Fig. 2d) were obtained from the Victorian Clinical Genetics Service. In brief, variants identified from clinical-grade targeted mtDNA sequencing were assessed using criteria adapted from the American College of Medical Genetics mitochondrial variant classification guidelines^12^, as described previously^64^. Variants were curated by medical genomics scientists, and variant classifications were reviewed by a multidisciplinary team. Curated missense variants are listed in Supplementary Dataset 3.

### Replication dataset

HelixMTdb population data was obtained as described above^30^. Linear models fitting neutral variation observed in HelixMTdb and their mutational likelihoods across loci were used to calculate expected values, as above. Variants at bases m.300-316, m.513-525, and m.16182-16194 were not called in HelixMTdb and therefore were excluded from calculations. Ratios of observed:expected variation for each functional class of mtDNA variation in HelixMTdb and their CIs were calculated as above.

### Regional constraint

A novel method was developed for assessment of mitochondrial regional constraint, which adapts methods described by Samocha *et al*^3^ and Davydov *et al*^65^ for nuclear constraint analyses. This analysis was implemented using custom scripts run in Python v3.10, as follows. For protein genes, the missense observed:expected ratio of all possible regions ≥30 bp within each gene was calculated, and a beta distribution used to compute the probability of the observed:expected ratio of each region being ≤ the gene’s missense observed:expected ratio. Regions with a p-value <0.01 were retained, and a greedy algorithm was applied to discard any region overlapping another with a lower p-value; for overlapping regions with the same p-value the longest was retained. This produced a list of non-overlapping candidate regions significantly more missense constrained than the gene. The false discovery rate (FDR) of each candidate was then estimated by applying the same method to 1000 random permutations of each gene, calculated as the proportion of permutations that produced a false positive result of the same length and ≤ p-value as the candidate region. Areas of regional missense constraint with FDR <0.1 were regarded as high-confidence and used for all analyses. Regional constraint in the rRNA genes was evaluated using the same process with minor modifications. All high-confidence regions are provided in Supplementary Dataset 2. A detailed description of this method is in the Supplementary Information.

The distance between residues and bases in 3D protein and rRNA structures was calculated using custom scripts implementing the Bio.PDB Biopython module^66^ in Python v3.10, to identify those in close proximity to regional constraint. The electron microscopy structures of human complex I (PDB:5XTD)^67^, complex III (PDB:5XTE)^67^, complex IV (PDB:5Z62)^68^, and the mitochondrial ribosome (PDB:6ZSE)^69^ from Protein Data Bank (PDB) were used. For the human complex V subunit MT-ATP6, the 3D structure predicted by AlphaFold obtained from UniProt was used (AF-P00846-F1)^70^. A protein residue was regarded to be in close proximity to regional constraint when the minimum distance between its alpha carbon atom and the alpha carbon of a residue in regional constraint was <6 Ångstrom, a threshold commonly used to define contacting residues^71^. A rRNA base was regarded to be in close proximity to regional constraint when the minimum distance between its nitrogen atom involved in base pairing and the equivalent nitrogen of a base in regional constraint was <6 Ångstrom, or if its phosphate atom and the phosphate of a base in regional constraint was <6 Ångstrom. Nitrogen atoms at position N1 were used for purines and position N3 for pyrimidines to capture base pair interactions, and phosphate atoms to capture flanking bases.

### Mitochondrial local constraint score

The mitochondrial local constraint (MLC) score was developed to measure local intolerance to base or amino acid substitutions at and around every position in the human mtDNA. This score was calculated using an overlapping sliding window method, implemented with custom scripts run in Python v3.10. Starting from position m.1, a window of length *k* was drawn and the observed:expected ratio of all of substitutions within the window and its 90% CI calculated. The window start position was moved by 1 bp, and the process repeated until all possible windows of length *k* in the mtDNA were evaluated. A window length *k* of 30 bp was used to enable all to have expected value >10. For positions in protein genes only amino acid substitutions (missense) were included in calculations, while all base substitutions were included for positions in RNA genes and non-coding regions. For each mtDNA position, the mean observed:expected ratio CI upper bound fraction (OEUF) of all *k* overlapping windows was computed and percentile ranked to achieve a score between 0.0 and 1.0 (ranging from least to most constrained). The score for every base position is provided in Supplementary Dataset 6. A MLC score was then assigned for every possible SNV as follows: non-coding, RNA and missense variants were assigned their positional score, and non-missense in protein genes were assigned scores based on the variant class OEUF with synonymous, stop gain, and start or stop lost assigned scores of 0.0, 1.0, and 0.70 respectively. A higher score is predicted to be more deleterious; scores for every SNV are in Supplementary Dataset 7. A detailed description of this method is in the Supplementary Information.

### Odds ratio enrichment analysis

Odds ratio (OR) analysis assessing pathogenic versus benign variation across categories of regional constraint and MLC score quartiles was calculated as 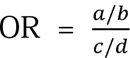, as previously described^2^, where *a* is the number of pathogenic variants in the category/quartile, *b* is the number of benign variants in the category/quartile, *c* is the number of pathogenic variants not in the category/quartile, and *d* is the number of benign variants not in the category/quartile. The standard error was calculated as 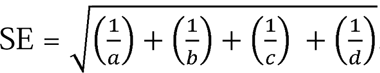. A 95% confidence interval for each OR was calculated from the SE, as *e^ln^*^[*O R*]±(1.96xSE)^. Pathogenic variants included those with a pathogenic or likely pathogenic classification in ClinVar or a confirmed disease association in MITOMAP. Benign variants were those with a benign classification in ClinVar.

### Visualization of protein and RNA 3D structures

Protein and rRNA 3D structures from Protein Data Bank (PDB) were visualized using UCSF ChimeraX^72^ v1.3. The electron microscopy structures of human complex I (PDB:5XTD)^67^, complex III (PDB:5XTE) ^67^, complex IV (PDB:5Z62)^68^, and the mitochondrial ribosome including A/P-site and P/E-site tRNAs (PDB:6ZSE)^69^ were used. The ovine complex I electron microscopy structure (PDB:6ZKM)^34^ was also used to show the homologous MT-ND4 region binding rotenone in Fig. 5a. Figures and videos displaying constraint data on 3D structures were generated using custom ChimeraX command files.

### Evaluation of heteroplasmy and blood cell counts in UK Biobank

Mitochondrial heteroplasmy was identified from whole genome sequencing (WGS) data from the UK Biobank, a large population study of people from the United Kingdom aged 40-69 years^73^, as described previously^49^. In brief, MitoHPC^74^ was used to call heteroplasmic SNVs with a heteroplasmy level of >5%. Variants were filtered using the following criteria: at poly-C homopolymer regions, read depth <300, and or with base quality, strandedness, slippage, weak evidence, germline, or position flags. Samples were excluded using the following criteria: mitochondrial contamination level >3%, two or more variants from a different mitochondrial haplogroup, multiple variants predicted as nuclear-encoded mitochondrial sequences, low coverage, mtDNA copy number ≤40, and or heteroplasmy count above five. Samples with cell count outliers more than three standard deviations from the mean were also excluded. 193,115 of 200,000 samples with WGS data were retained for analysis. The association between heteroplasmy metrics (count and MLC score sum) and cell counts (platelets and neutrophils) was determined using a linear regression model adjusting for age (natural spline, 4 degrees of freedom), sex, and smoking status (“current”, “former”, or “never smoker”). Raw neutrophil count was transformed using log(neutrophil count + 1) to more closely approximate a normal distribution. Regression models were run with each metric separately (‘Single’) or together in the same model (‘Combined’) in R v4.0.4.

### Statistical analysis

The statistical tests utilized in this study are described in detail in the relevant Methods and or Supplementary Methods sections.

### Data availability

Data analyzed or generated during this study are included in this published article and its Supplementary files, and or available via https://github.com/leklab/mitochondrial_constraint.

### Code availability

The code used to perform all analyses and generate all figures in this manuscript are available at https://github.com/leklab/mitochondrial_constraint.

## Acknowledgements

N.J.L. received a National Health and Medical Research Council (NHMRC) Early Career Fellowship APP1159456 and an Australian American Association Scholarship. This research was conducted using the UK Biobank Resource under Application Number 17731, and supported by National Heart, Lung and Blood Institute, National Institutes of Health (NIH) grant R01HL144569. The content is solely the responsibility of the authors and does not necessarily represent the official views of the NIH. D.R.T. was supported by an NHMRC Principal Research Fellowship GNT1155244. Research conducted at the Murdoch Children’s Research Institute was supported by the Victorian Government’s Operational Infrastructure Support Program. We are grateful to all of the families in SPARK, the SPARK clinical sites and SPARK staff, and we appreciate obtaining access to the data on SFARI Base. We gratefully acknowledge Sarah Calvo for providing advice on the constraint model and its application to gnomAD. We also acknowledge the contributions of the broader team involved in curation of disease-associated variation at the Victorian Clinical Genetics Service, including Belinda Chong and Sebastian Lunke.

## Author contributions

N.J.L. and M.L. contributed study conception and design. N.J.L., S.L.B., K.M.L., G.T., D.P., A.G.C., S.C., J.C., D.R.T., D.E.A., and M.L. contributed to data acquisition. N.J.L., W.L., H.Z., S.R.S, and M.L. contributed to the methods development of the model. N.J.L., W.L., S.L.B., D.E.A., and M.L. contributed to data analysis. N.J.L., W.L., S.L.B., K.M.L., G.T., A.G.C., S.C., J.C., D.R.T., H.Z., D.E.A., S.R.S, and M.L. contributed to interpretation of the data. M.L. supervised and managed the study. N.J.L. and M.L. drafted the manuscript. All authors contributed to manuscript review and editing.

## Additional Information

Supplementary Information is available for this paper. This contains Supplementary Methods, Supplementary Figures 1-5 and Supplementary Tables 1-2 which pertain to the Supplementary Methods, Supplementary Discussion, Supplementary References, and descriptions of Supplementary Datasets and the Supplementary Video.

Correspondence and requests for materials should be addressed to Nicole J. Lake or Monkol Lek.

## Extended Data Figures

**Extended Data Figure 1:**
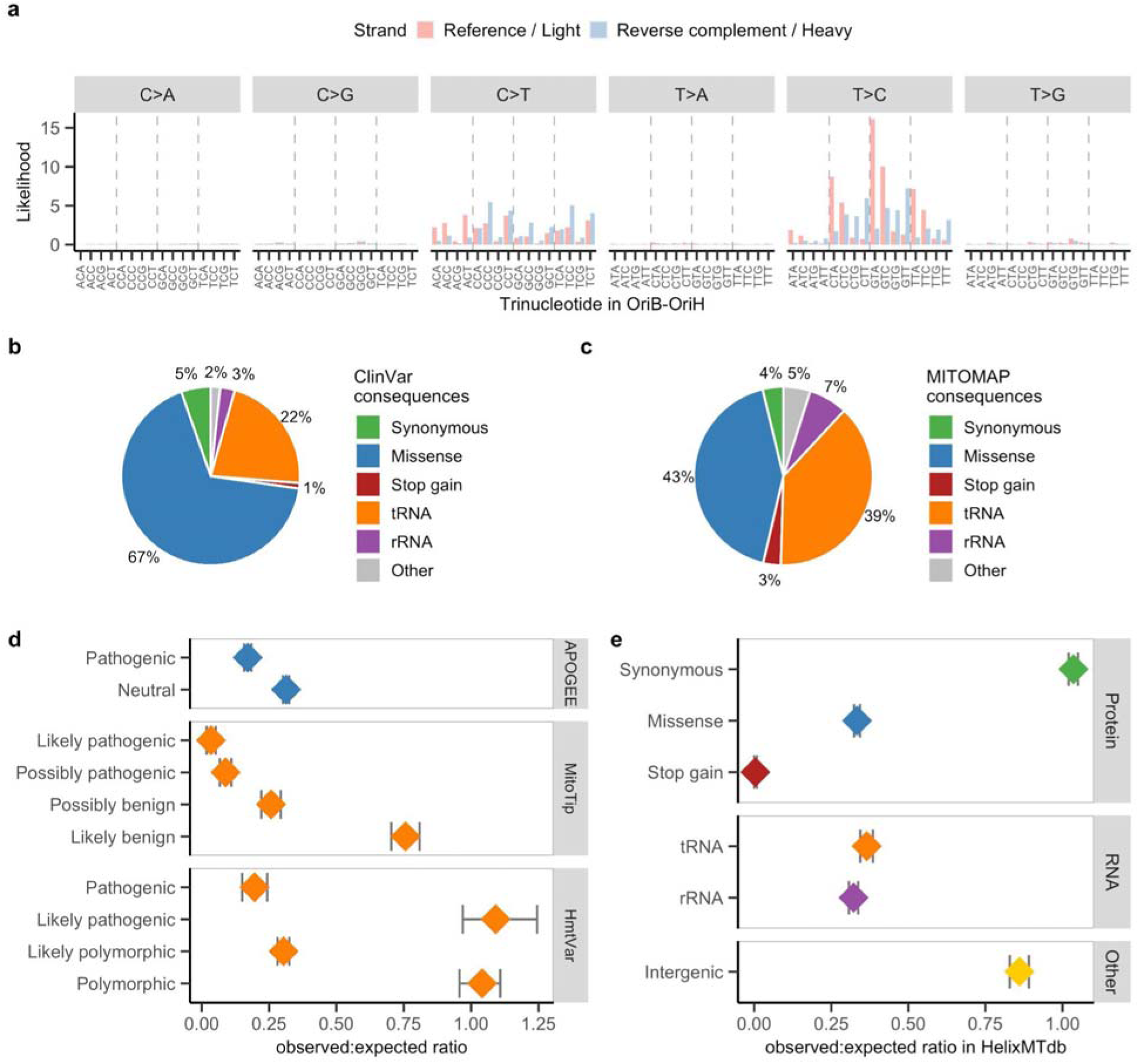
Mutability, disease-associated variation, and constraint across classes of human mtDNA variation. **(a)** Trinucleotide mutational signature of mtDNA mutations within the OriB-OriH region (m.16197-191) predicted by the composite likelihood model. Mutation likelihoods for the six pyrimidine base substitution types across 96 trinucleotides are shown, colored by whether the reference nucleotide is in the reference ‘light’ or reverse complement ‘heavy’ strand. **(b-c)** Proportion of total disease-associated variants in ClinVar (*n*=2607) **(b)** and MITOMAP (*n*=882) **(c)** by consequence. **(d)** The observed:expected ratio of *in silico* predictions in gnomAD for missense variants by APOGEE (pathogenic, *n*=7276 and neutral, *n*=16,800), and for tRNA variants by MitoTIP (likely pathogenic, *n*=981; possibly pathogenic, *n*=1171, possibly benign, *n*=1162 and likely benign, *n*=1198) and HmtVar (pathogenic, *n*=202; likely pathogenic, *n*=6, likely polymorphic, *n*=4139 and polymorphic, *n*=24); all of which are recommended per ACMG/AMP mtDNA guidelines for variant interpretation. Note the outlier HmtVar ‘likely pathogenic’ group only includes six variants. **(e)** Assessment of functional classes of mtDNA variation in a replication dataset, HelixMTdb. The *n* per class is per Fig. 1d. Bars in (d-e) represent 90% confidence intervals.

**Extended Data Figure 2:**
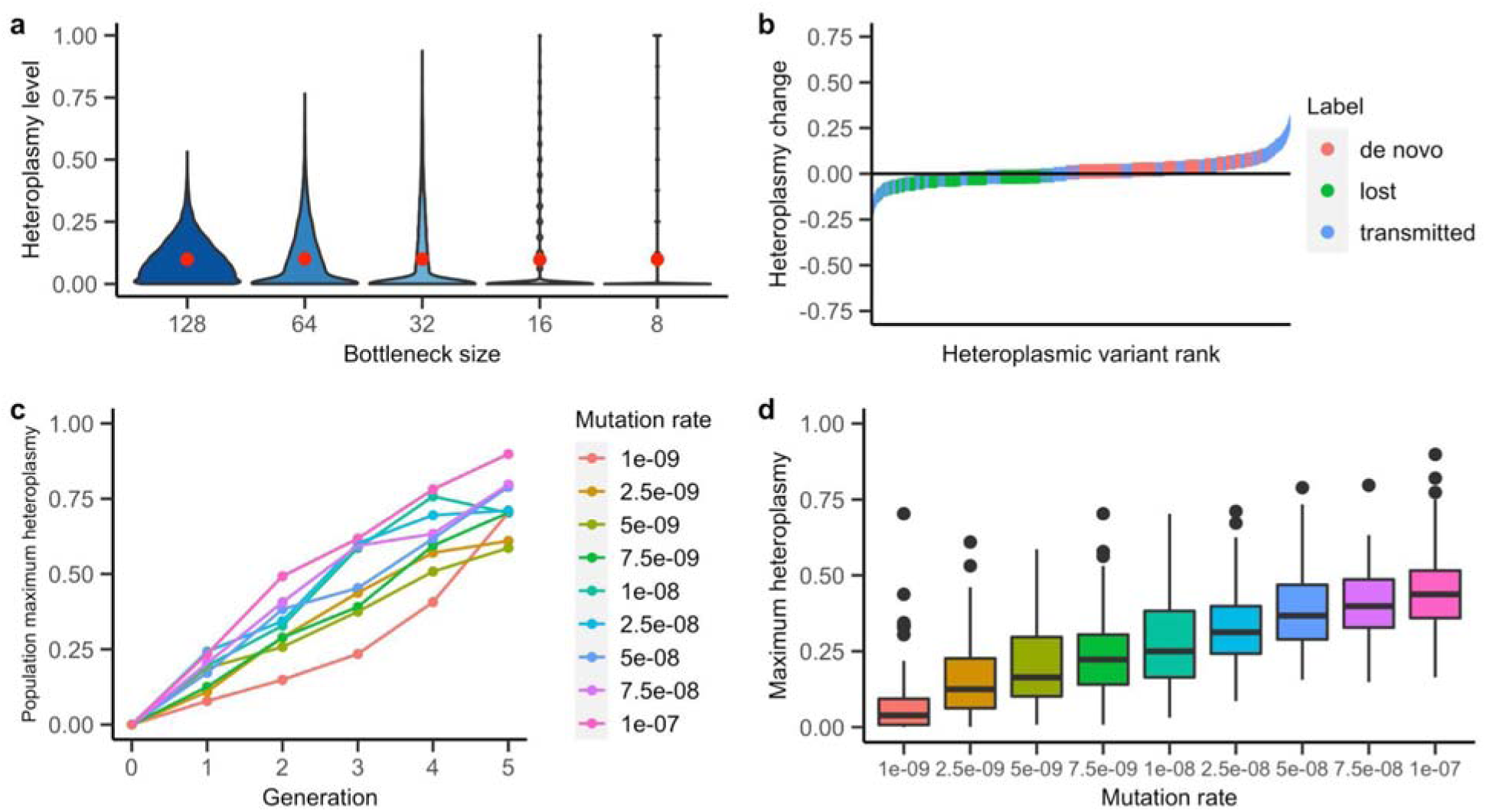
Simulation of germline mtDNA mutation and heteroplasmy in maternal lineages. **(a)** The simulated heteroplasmy distribution of mutations in 10,000 offspring who inherited a mutation with heteroplasmy 0.1, for five different bottleneck sizes; the mean is shown in red. This plot reproduces an analysis by Colnaghi *et al*^59^, using a mutation rate of 10^-8^. For (b-d) we used their standard bottleneck size of 128. **(b)** Heteroplasmy changes between offspring (generation 5) and mothers (generation 4) across 10,000 lineages, ordered by degree of shift. *De novo* represents variants with heteroplasmy >0.01 in offspring and <0.01 in mothers, lost have heteroplasmy <0.01 in offspring and >0.01 in mothers, and transmitted have heteroplasmy >0.01 in offspring and mothers, to match criteria used by Wei *et al*^19^. A mutation rate of 10^-8^ was used; starting heteroplasmy was 0 for generation 0. **(c)** Maximum heteroplasmy across 10,000 lineages across five generations, colored by mutation rate. **(d)** Boxplots showing maximum heteroplasmy distribution from 1000 bootstrap replicates sampling 100 individuals from the population (*n*=10,000) at generation five; starting heteroplasmy was 0 for generation 0.

**Extended Data Figure 3:**
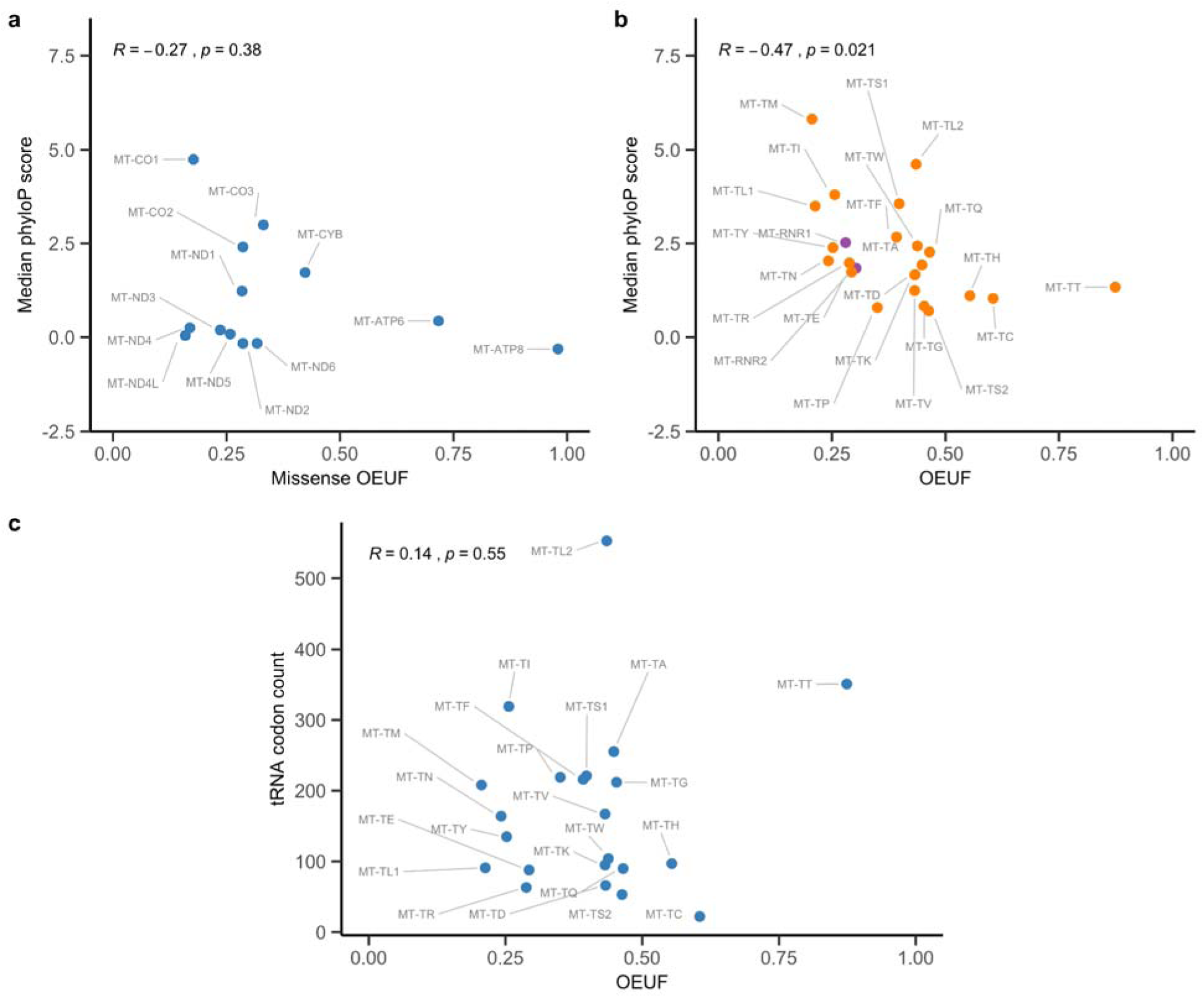
Assessment of conservation and codon usage across genes. Median phyloP base conservation scores for each protein **(a)** or RNA **(b)** gene, derived from 100 vertebrate genomes. Higher phyloP values represent increased conservation. **(c)** The count of codons within the protein-coding sequence corresponding to each tRNA. Pearson correlation coefficient (R) and its p-value (p) is shown in (a-c). OEUF stands for observed:expected ratio confidence interval upper bound fraction, and OEUF values for (a-c) are provided in Supplementary Dataset 1.

**Extended Data Figure 4:**
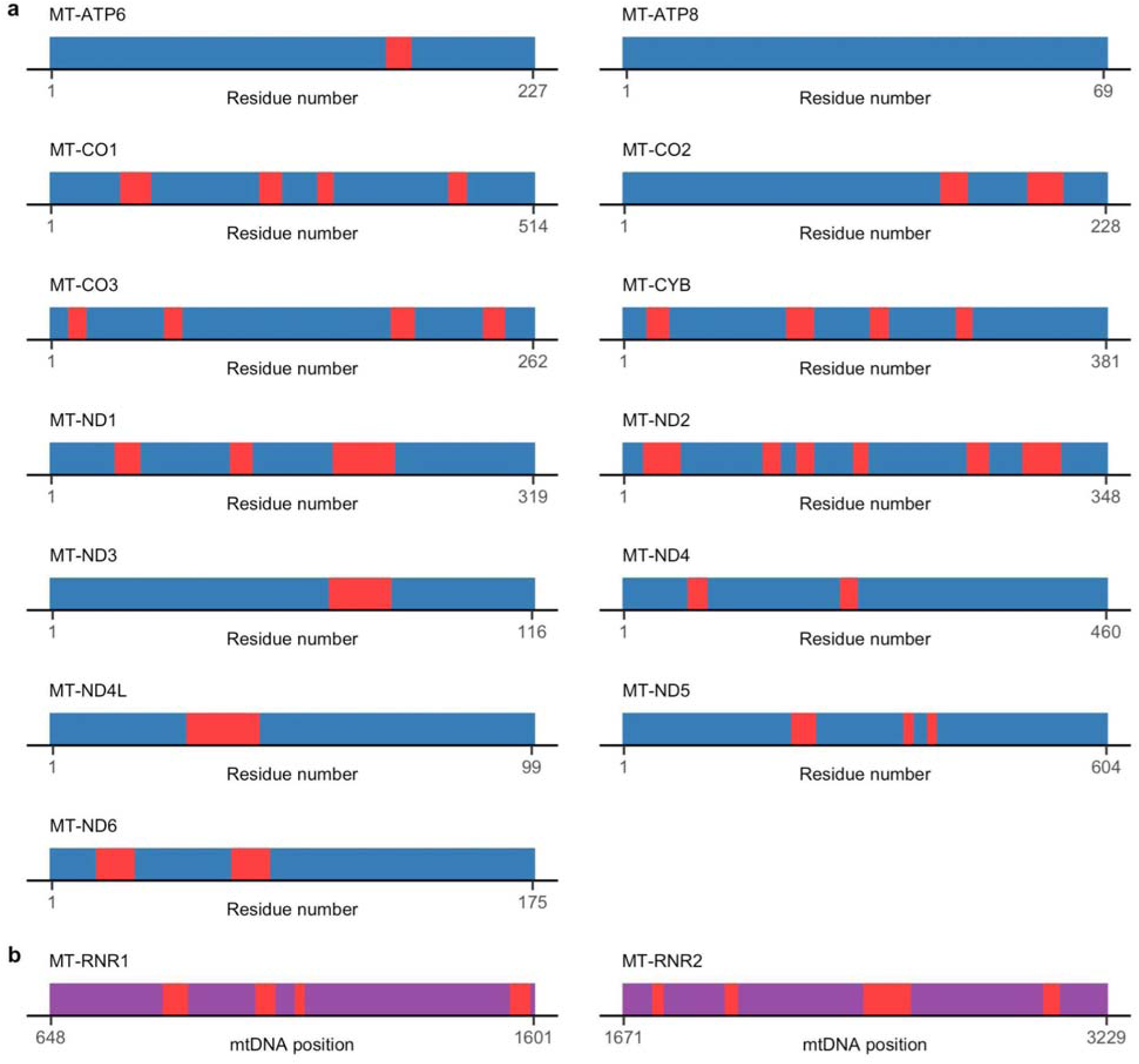
Areas of regional constraint within each protein and rRNA gene. **(a)** Intervals of regional missense constraint identified in each protein are colored in red. For display purposes each protein is shown at the same length (i.e. are not scaled by their actual protein length), and amino acid residue numbering is shown. **(b)** Intervals of regional constraint identified in each rRNA are colored in red. The rRNA sequences are not scaled by their length, and mtDNA position coordinates are shown. Coordinates for (a-b) are provided in Supplementary Dataset 2.

**Extended Data Figure 5:**
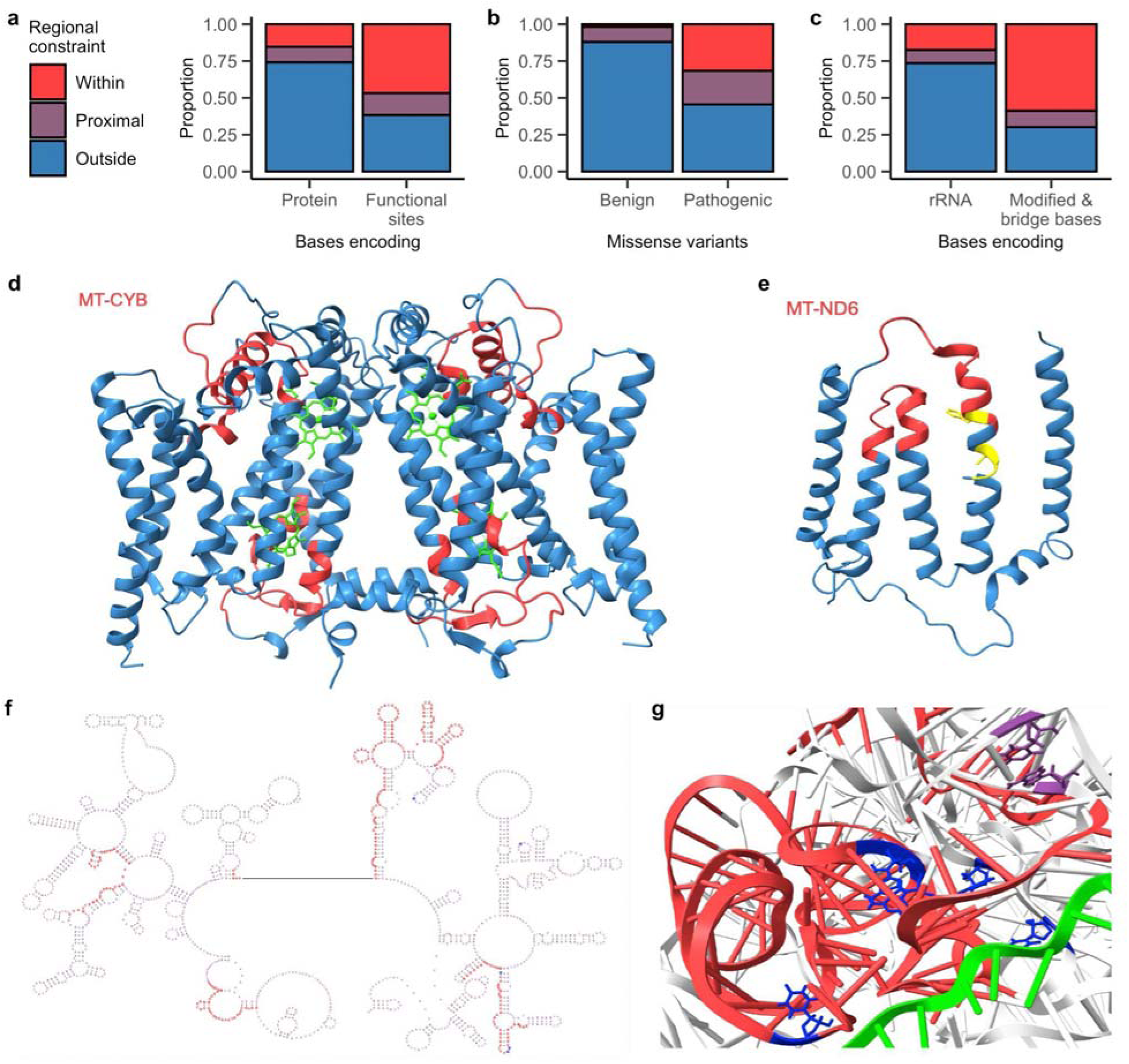
Characteristics of regional constraint. **(a-b)** The proportion of bases encoding proteins (*n*=11,341) or residues of functional significance (*n*=141) **(a)** or benign (*n*=625) and pathogenic (*n*=79) missense variants (most severe consequence) **(b)** that are within, proximal to (<6 Ångstrom distance from), or outside regional missense constraint. **(c)** The proportion of bases encoding rRNA (*n*=2512), or modified bases and bases in rRNA:rRNA intersubunit bridges (*n*=63) that are within, proximal to (<6 Ångstrom distance from), or outside regional constraint. **(d)** The four areas of regional missense constraint in MT-CYB are shown in red, visualized in its dimeric 3D structure. Heme molecules involved in electron transfer are colored green. **(e)** The two areas of regional missense constraint in MT-ND6 are shown in red in the 3D structure. Residues colored in yellow are involved in the transition from the open to closed complex state in the π-bulge (p.61-63 and p.67) per Kampjut and Sazanov^34^. **(f)** The areas of regional constraint within the MT-RNR2 secondary structure, indicated by red font; modified bases are in blue font. **(g)** An area of regional constraint within the MT-RNR1 tertiary structure, indicated in red. Modified bases are colored blue, and disease-associated bases (m.1494 and m.1555) purple. The mRNA molecule is colored green.

**Extended Data Figure 6:**
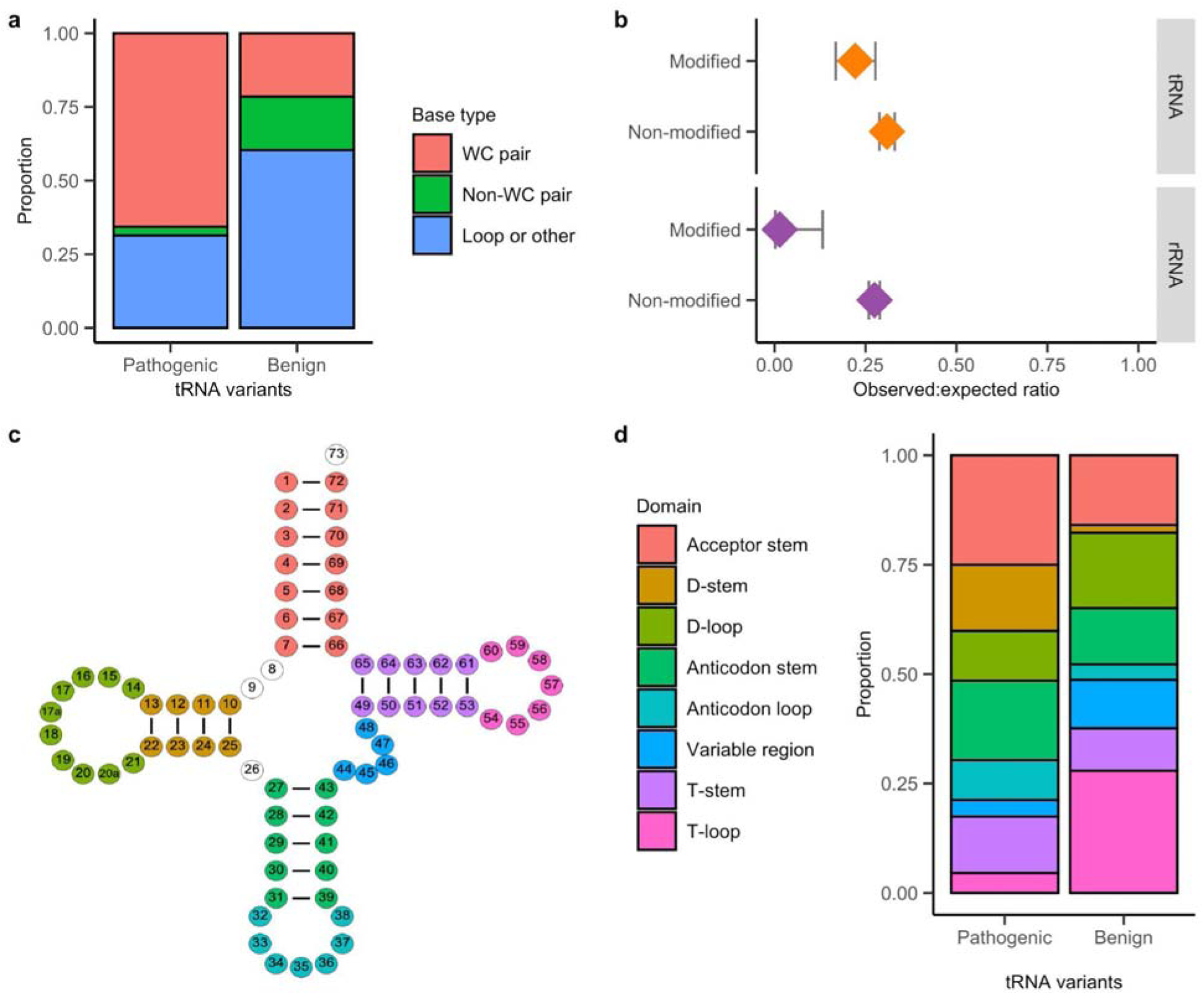
Characteristics of RNA variants and bases. **(a)** The proportion of pathogenic (*n*=121) and benign (*n*=232) tRNA variants for each base type. **(b)** The observed:expected ratio for variants in modified and non-modified bases in tRNA (modified, *n*=411 and non-modified, *n*=4101) and rRNA (modified, *n*=30 and non-modified, *n*=7506). Error bars represent the 90% confidence interval. **(c)** The generic tRNA secondary structure, with positions colored by domain. **(d)** The proportion of pathogenic (*n*=121) and benign (*n*=232) tRNA variants for each domain, following the color legend in (c).

**Extended Data Figure 7:**
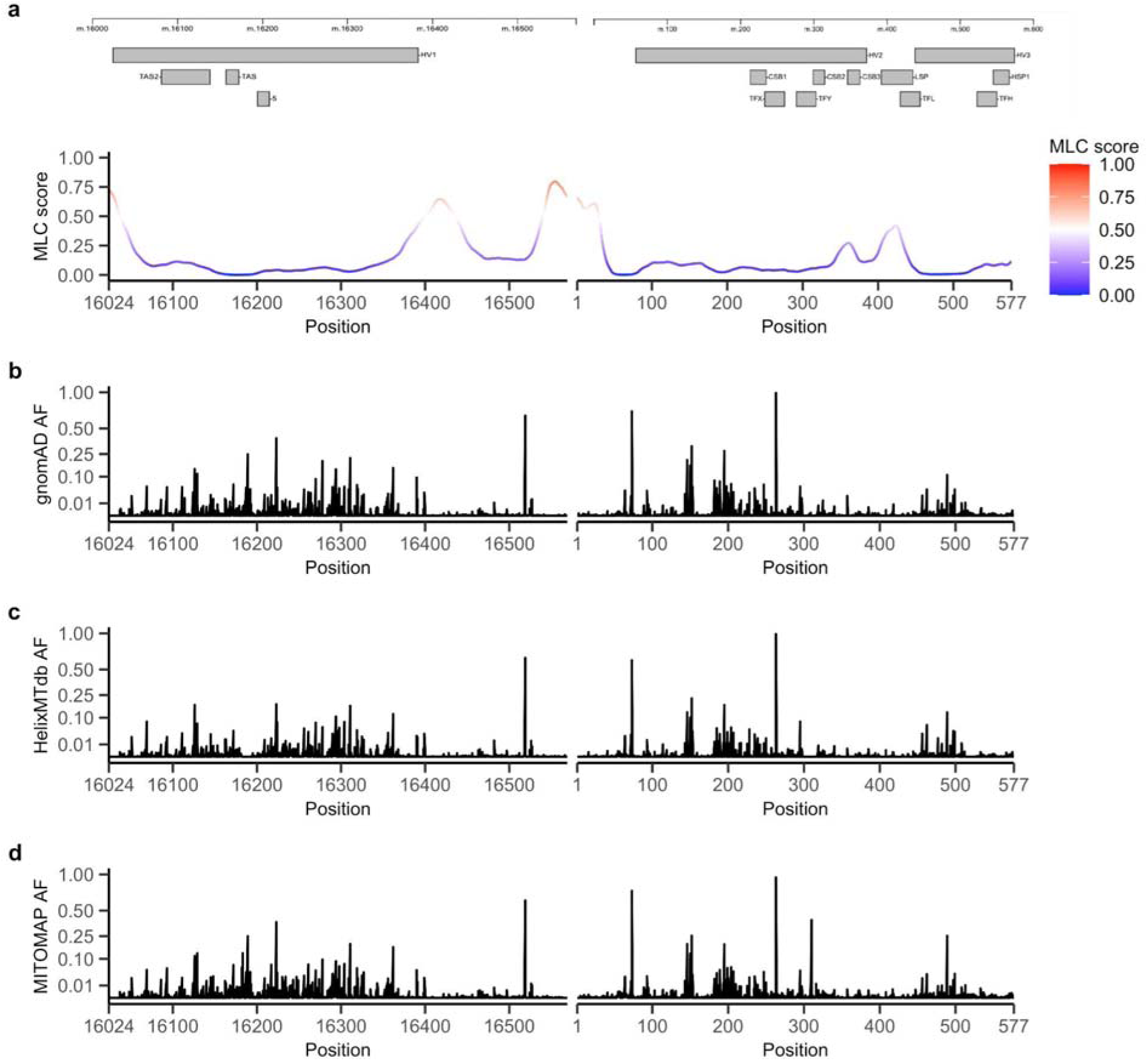
Mitochondrial local constraint (MLC) scores and population allele frequencies across the non-coding control region. **(a)** The MLC score of positions across the control region are shown; a schematic of annotated non-coding elements is displayed above. The five peaks from left to right overlap (1) a recently discovered second light strand promoter^47^, (2-3) regions of unknown function within the D-loop, (4) conserved sequence block 3, or (5) the light strand promoter. Base scores are provided in Supplementary Dataset 6. **(b-c)** The homoplasmic allele frequency (AF) of variants across the control region in gnomAD (b) or HelixMTdb (c). **(d)** The allele frequency of variants across the control region in the MITOMAP database. (b-d) are displayed with a square root transformed y-axis; note only SNVs are included.

**Extended Data Figure 8:**
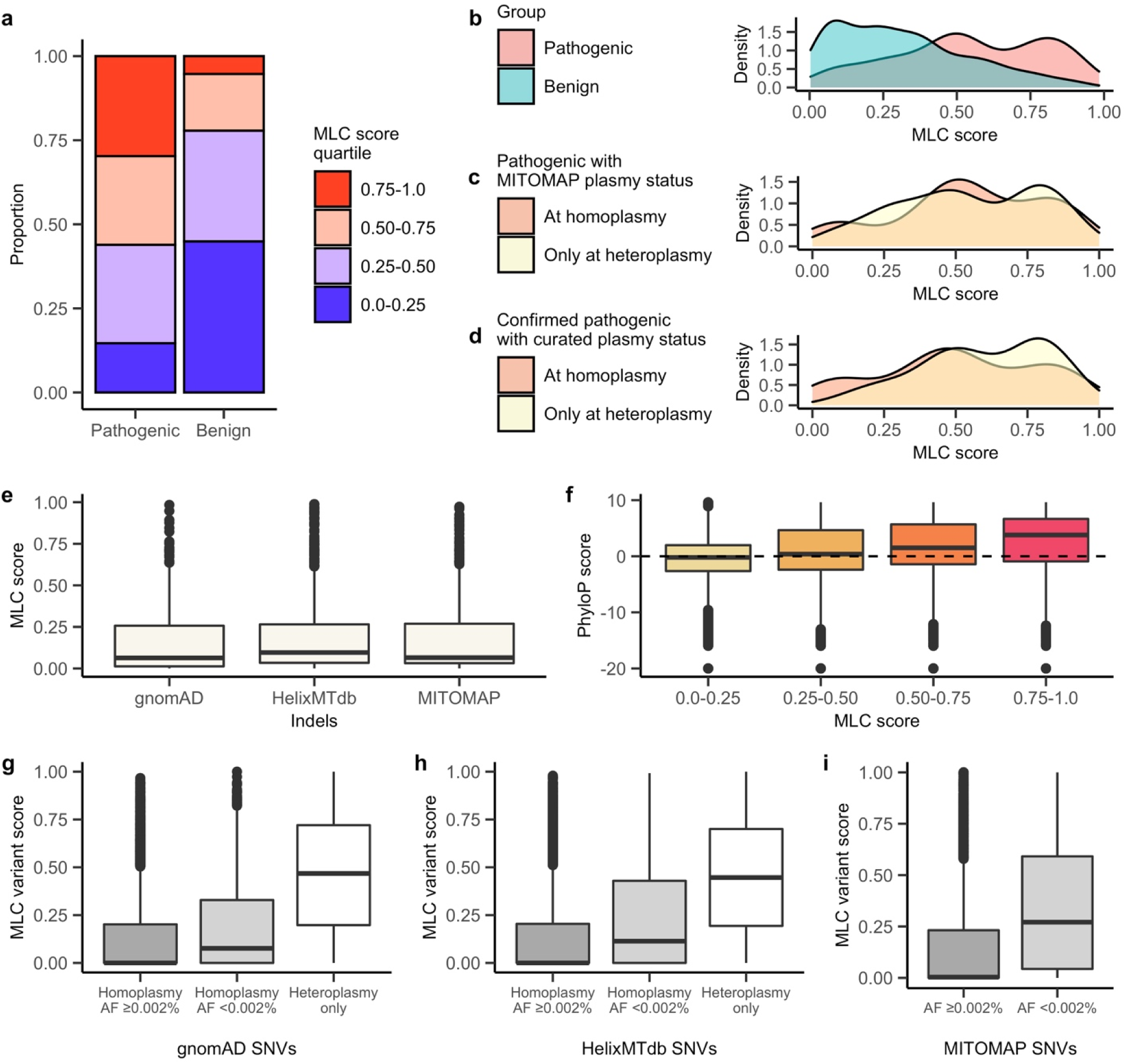
Relationship between the mitochondrial local constraint (MLC) score and genomic annotations. **(a)** The proportion of benign (*n*=884) and pathogenic (*n*=205) variants in each score quartile. **(b)** Density plot showing the score distribution of disease-associated variants; numbers per (a). **(c)** Density plot showing the score distribution of 184 pathogenic variants with disease plasmy status in MITOMAP, colored by association with disease at heteroplasmy only, or at homoplasmy. **(d)** Density plot showing the score distribution of 88 ‘confirmed’ pathogenic variants from MITOMAP, colored by whether reported in individuals at heteroplasmy only or at homoplasmy, per a manual literature review. Plots (a-d) include missense and RNA variants only, and for (c-d) ‘at homplasmy’ includes observed at both homoplasmy and heteroplasmy. **(e)** Boxplot showing the score distribution for base positions where indels are observed in gnomAD (*n*=416), HelixMTdb (*n*=697), and MITOMAP (*n*=667) databases. **(f)** The distribution of PhyloP base conservation scores for bases within each score quartile. PhyloP scores >0 (dashed horizontal line) indicate conserved sites by PhyloP. **(g-i)** The MLC variant score distribution for SNVs across population frequency categories in gnomAD (homoplasmy AF ≥0.002%, *n*=7363; homoplasmy AF <0.002%, *n*=1846 and heteroplasmy only, *n*=1641) (g), HelixMTdb (homoplasmy AF ≥0.002%, *n*=8049; homoplasmy AF <0.002%, *n*=3442 and heteroplasmy only, *n*=2613) (h) and MITOMAP (AF ≥0.002%, *n*=8617 and AF <0.002%, *n*=10,343) (i) databases. Note that allele frequency <0.002% is recommended as evidence of pathogenicity in ACMG/AMP mtDNA guidelines^12^, and that heteroplasmy data is not available for MITOMAP.

