## Supplementary Information for "Quantifying constraint in the human mitochondrial genome"

### Supplementary Methods

#### 1. Mitochondrial composite likelihood model

We adapted the composite likelihood model described by Dietlein et al to quantify mutability in the human mitochondrial DNA (mtDNA)[1]. This model was developed for the study of passenger mutations in cancer nuclear exomes, which can have a sparsity of mutation counts per possible nucleotide context[1]. This approach was therefore well suited to handle the possible sparsity of counts per context associated with the smaller genome size of the mtDNA. We made several adaptations to the composite likelihood model to enable its application to the mtDNA. These modifications aimed to handle the known replicative strand bias for transitions[2], unequal frequency of pyrimidines and purines in the reference sequence[3], as well as the inverted signature for transitions within the non-coding OriB-OriH region[2].

We quantified mutability using de novo mutations ascertained from the literature and an in-house dataset (described below). Since the mutability of a mtDNA base can depend on the sequence context at the +1 and -1 positions[2,4], we quantified mutability in trinucleotide contexts. The composite model decomposes the mutational likelihood of each trinucleotide context into multiplicative factors, namely the effects of the base substitution type $\lambda_t$, the flanking 5' nucleotide $\lambda_{-1}$ and the flanking 3' nucleotide $\lambda_{+1}$. The likelihood of the base substitution type $\lambda_t$ can be further decomposed into the effect of the reference nucleotide $\lambda_{\boldsymbol{n(t)}}$ and the mutation class $\lambda_{\boldsymbol{c(t)}}$. A mutation likelihood score $\lambda$ is obtained for every possible base substitution at every nucleotide in the mtDNA, where $\lambda > 1$ indicates increased likelihood of mutation, and $\lambda < 1$ a decreased likelihood compared with genome-wide average. Our mitochondrial composite likelihood model is described in detail below.

#### 1.1 Mutation classification system and reference sequence

We classified the six pyrimidine base substitution types by whether the reference nucleotide C or T is on the reference or reverse complement strand (i.e. G and A in the reference strand). This classification of 12 base substitution types $t$ and their reference nucleotide $n(t)$ is summarized in Supplementary Table 1. These mutations are further categorized into three mutational classes $c(t)$: transversions type I (class I), transversions type II (class II) and transitions (class III).

We used the GRCh38 mitochondrial reference sequence NCBI accession NC_012920.1, also known as the revised Cambridge Reference Sequence (rCRS). Due to the different mutational signature within the non-coding OriB-OriH region spanning m.16197-16569 and m.1-191 (across the artificial break)[2], we computed mutability within the reference sequence excluding OriB-OriH (m.192-16196) separately to the OriB-OriH region.

**Supplementary Table 1: Mutation classification system used.** This includes 12 mutation types, and three mutation classes. The pyrimidine mutation types are also shown.

| **Mutation** | **Reference nucleotide $n(t)$** | **Mutation type $t$** | **Mutation class $c(t)$** | **Pyrimidine mutation type $pyr(t)$** | **Pyrimidine reference nucleotide $pyr(n(t))$** |
|---|---|---|---|---|---|
| C>A | C | 1 | I | C>A | C |
| C>G | C | 2 | II | C>G | C |
| C>T | C | 3 | III | C>T | C |
| T>A | T | 4 | I | T>A | T |
| T>C | T | 5 | III | T>C | T |
| T>G | T | 6 | II | T>G | T |
| G>T | G | 7 | I | C>A | C |
| G>C | G | 8 | II | C>G | C |
| G>A | G | 9 | III | C>T | C |
| A>T | A | 10 | I | T>A | T |
| A>G | A | 11 | III | T>C | T |
| A>C | A | 12 | II | T>G | T |

#### 1.2 Likelihood ratio for mutation at reference nucleotides

We counted the number of de novo mutations of each base substitution type $t$ and summarized them into type count vector $\boldsymbol{v^{type}}$. Each element $v_t^{type}$ corresponds to the number of base substitutions of type $t \in \{1, \dots, 12\}$. We use $v_t^{type}$ to define the likelihood ratios for base substitution at the reference nucleotides:

$$\lambda_C = \frac{\left(v_1^{type} + v_2^{type} + v_3^{type}\right)}{|v^{type}| \cdot P(n(t)=C)}$$

$$\lambda_T = \frac{\left(v_4^{type} + v_5^{type} + v_6^{type}\right)}{|v^{type}| \cdot P(n(t)=T)}$$

$$\lambda_G = \frac{\left(v_7^{type} + v_8^{type} + v_9^{type}\right)}{|v^{type}| \cdot P(n(t)=G)}$$

$$\lambda_A = \frac{\left(v_{10}^{type} + v_{11}^{type} + v_{12}^{type}\right)}{|v^{type}| \cdot P(n(t)=A)}$$

where $|v^{type}| := \sum_t v_t^{type}$ (sum of all mutations across all 12 types), and $P\big(n(t)\big)$ reflects our a priori assumption to observe mutations at each $n(t) \in \{A, C, G, T\}$ at the same frequency as the $n(t)$ within the reference sequence (i.e. the proportion of the $n(t)$ in the reference sequence).

#### 1.3 Likelihood ratio for the mutation classes

We used $v_t^{type}$ to define the likelihood ratios for the mutation classes I, II, III:

$$\lambda_I = \frac{\left(v_1^{type} + v_4^{type} + v_7^{type} + v_{10}^{type}\right)}{|v^{type}| \cdot P(c(t)=I)}$$

$$\lambda_{II} = \frac{\left(v_2^{type} + v_6^{type} + v_8^{type} + v_{12}^{type}\right)}{|v^{type}| \cdot P(c(t)=II)}$$

$$\lambda_{III} = \frac{\left(v_3^{type} + v_5^{type} + v_9^{type} + v_{11}^{type}\right)}{|v^{type}| \cdot P(c(t)=III)}$$

where $|v^{type}| := \sum_t v_t^{type}$, and $P\big(c(t)\big)$ is the probability of the mutation class, calculated as the sum of probabilities for each mutation type $t$ in the class where

$$c(t) = \begin{cases} I & t = \mathrm{C > A, T > A, G > T, A > T} \\ II & t = \mathrm{C > G, T > G, G > C, A > C} \\ III & t = \mathrm{C > T, T > C, G > A, A > G} \end{cases}$$

The probability of each type $t$ is estimated using the relative frequency of nucleotides in the reference sequence. Specifically, we calculate this as the frequency of the reference nucleotide $n(t)$ multiplied by the frequency of the alternate nucleotide amongst all possible alternates in the reference sequence, i.e. $P(C > A) = P(n(t) = C) \cdot (P(n(t) = A)/(1 - P(n(t) = C))$.

#### 1.4 Likelihood of mutation type

We used the likelihood ratios $\lambda_{n(t)}$ for the reference nucleotide and $\lambda_{c(t)}$ for the mutation class to calculate the likelihood ratio of each base substitution type

$$\lambda_t = \lambda_{n(t)} \cdot \lambda_{c(t)}$$

for each $n(t)$ reference nucleotide and $c(t)$ mutation class defined in Supplementary Table 1.

#### 1.5 Likelihood ratio for sequence context for class III mutations

We use $v_t^{type}$, $v_{t,p,n}^{seq}$ and $f_{n(t),p,n'}^{ref}$ (defined below) to determine the likelihood ratios for sequence context around class III mutations:

$$\lambda_{t,p,n} = \frac{v_{t,p,n}^{seq}}{v_t^{type} \cdot f_{n(t),p,n'}^{ref}}$$

where $v_{t,p,n}^{seq}$ is the count of nucleotide $n \in \{A, C, G, T\}$ at position $p \in [-1:1] \backslash \{0\}$ around base substitution of type $t \in \{\mathrm{C > T, T > C, G > A, A > G}\}$, $v_t^{type}$ is the count of base substitutions of type $t$, and $f_{n(t),p,n'}^{ref}$ is the frequency of nucleotide $n' \in \{A, C, G, T\}$ at position $p \in [-1:1] \backslash \{0\}$ around the reference nucleotide $n(t)$. Specifically, $f^{ref}$ for class III mutations is calculated as:

$$f_{n(t),p,n'}^{ref} = \frac{f_{(n_p=n',n(t))}^{ref}}{P(n(t))}$$

where $P(n(t))$ is the proportion of the $n(t)$ in the reference sequence, and $f_{(n_p=n',n(t))}^{ref}$ is the frequency of observing nucleotide $n'$ at position $p$ around $n(t)$ across the total number of nucleotide sequences of length $(n_p, n(t))$ in the reference sequence.

#### 1.6 Likelihood ratio for sequence context for class I and class II mutations

For class I and II mutations (transversions), due to their lower counts we calculate $v^{seq}$ and $f_{n(t),p,n'}^{ref}$ by annotating them by their pyrimidine reference nucleotide (C or T) and calculating in a strand agnostic manner. While there is replicative strand bias for transitions, this has not been established for transversions[2,5], supporting this approach. Therefore, we classify the eight transversion mutation types $\boldsymbol{t}$ into their four pyrimidine mutation types $\boldsymbol{pyr(t)} \in \{\mathrm{C > A, C > G, T > A, T > G}\}$ (as per Supplementary Table 1).

We prepared $\boldsymbol{v_{pyr(t)}^{type}}$ where each element corresponds to the number of the $\boldsymbol{pyr(t)}$. We used $v_{pyr(t)}^{type}$ as well as $\boldsymbol{v_{pyr(t),p,n}^{seq}}$ and $f_{pyr(n(t)),p,n'}^{ref}$ (defined below) to determine the following likelihood ratios for the sequence context around class I and II mutations:

$$\lambda_{t,p,n} = \frac{v^{seq}_{pyr(t),p,n}}{v^{type}_{pyr(t)} \cdot f^{ref}_{pyr(n(t)),p,n'}}$$

where $\boldsymbol{pyr(t)}$ is the pyrimidine mutation type of the mutation type *t*, $v^{type}_{pyr(t)}$ is the count of the $\boldsymbol{pyr(t)}$, and $v^{seq}_{pyr(t),p,n}$ is the count of nucleotide $n \in \{A, C, G, T\}$ at position $p \in [-1{:}\,1]\{0\}$ around $\boldsymbol{pyr(t)}$. For mutation types $t = \mathrm{G > T, G > C, A > T, A > C}$ where the reference nucleotide $n(t)$ is A or G, the complement $\bar{n}$ of the flanking nucleotide $n$ at position $-p$ is counted instead. Therefore $v^{seq}_{pyr(t),p,n}$ is calculated as:

$$v^{seq}_{pyr(t),p,n} = \begin{cases} v^{seq}_{pyr(t),p,n} & \text{for } t = \mathrm{C > A, C > G, T > A, T > G} \text{ where } n(t) = \mathrm{C, T} \\ v^{seq}_{pyr(t),-p,\bar{n}} & \text{for } t = \mathrm{G > T, G > C, A > T, A > C} \text{ where } n(t) = \mathrm{A, G} \end{cases}$$

$f^{ref}_{pyr(n(t)),p,n'}$ is the frequency of nucleotide $n' \in \{A, C, G, T\}$ at position $p \in [-1{:}\,1]\backslash\{0\}$ around the pyrimidine reference nucleotide $\boldsymbol{pyr}(n(\boldsymbol{t}))$, where:

$$pyr\big(n(t)\big) = \begin{cases} C & n(t) = C, G \\ T & n(t) = T, A \end{cases}$$

For reference nucleotides $n(t)$ that are A or G, the frequency of the complement $\overline{n'}$ of the flanking nucleotide $n$ at position $-p$ is counted instead. Specifically, $f^{ref}_{pyr(n(t)),p,n'}$ is calculated as:

$$f^{ref}_{pyr(n(t)),p,n'} = \begin{cases} \dfrac{f^{ref}_{(n_p = n', pyr(n(t)))}}{P(pyr(n(t)))} & where\ n(t) = C, T \\ \dfrac{f^{ref}_{(n_{-p} = \overline{n'}, pyr(n(t)))}}{P(pyr(n(t)))} & where\ n(t) = A, G \end{cases}$$

where $P(pyr(n(t)))$ is the proportion of the $\boldsymbol{pyr}(n(\boldsymbol{t}))$ in the reference sequence, and $f^{ref}_{(n_p = n', pyr(n(t)))}$ is the frequency of observing nucleotide $n'$ at position $p$ around $\boldsymbol{pyr}(n(\boldsymbol{t}))$ across the total number of nucleotide sequences of length $(n_p, \boldsymbol{pyr}(n(\boldsymbol{t})))$ in reference sequence.

#### 1.7 Composite likelihood of position mutability based on sequence context

We used the likelihood ratios for sequence context $\lambda_{t,p,n}$ to define the composite likelihood of the flanking nucleotide sequence on the mutation probability:

$$\lambda_{t,(n_{-1},\ldots,n_1)} = \prod_{-1 \le p \le 1\ p \ne 0} \lambda_{t,p,n_p}$$

where the effect of the flanking sequence $(n_{-1}, \ldots, n_1) \in \{A, C, G, T\}^{1+1+1}$ of length *1+1+1* on base substitutions of type $t$ is modeled as a product of the effect of each flanking base, and $p$ = -1 represents the 5' flanking nucleotide while $p$ = 1 represents the 3' flanking nucleotide.

#### 1.8 Composite likelihood of each mutation type at each position

We used the likelihood ratio for the mutation type and the composite likelihood of sequence context to determine mutability of every possible base substitution within the reference sequence, first excluding the OriB-OriH region. Specifically, we compute the mutability of each mutation class $c$ at each mtDNA coordinate $m \in$ [192:16196] with reference nucleotide $n_0$ and flanking nucleotide $n_p$ at position $p$ as:

$$\lambda_{m,c} = \lambda_{n_0} \cdot \lambda_c \cdot \prod_{-1 \le p \le 1\ p \ne 0} \lambda_{t,p,n_p}$$

The replicative strand bias for transitions has been shown to be inverted in a segment of the non-coding control region bounded by the OriB site at m.16197 and the OriH site at m.191, which we refer to as the 'OriB-OriH' region[2]. Therefore we quantified mutability in this region separately. For determining the likelihood of mutation type, we only use de novo mutations and reference nucleotides within m.16197-191 to build $\boldsymbol{v^{type}}$, $\lambda_{n_0}^{ori}$ and $\lambda_c^{ori}$. For determining the likelihood of sequence context, we handle transitions and transversions separately. For transitions, we build $\boldsymbol{v^{seq}}$, $\boldsymbol{f^{ref}}$ and $\lambda_{t,p,n_p}^{ori}$ using de novo mutations and reference nucleotides within m.16197-191. For transversions, due to their lower counts we apply the likelihood of sequence context calculated previously ($\lambda_{t,p,n_p}$) for m.192-16196, on the assumption that sequence context around transversions is similar within the OriB-OriH, given the distinct mutational signature reported is for transitions only. This can be summarized as the following for $m \in$ [1-191, 16196-16569]:

$$\lambda_{m,c}^{ori} = \begin{cases} \lambda_{n_0}^{ori} \cdot \lambda_c^{ori} \cdot \prod_{-1 \le p \le 1\ p \ne 0} \lambda_{t,p,n_p} & where\ c(t) = I, II \\ \lambda_{n_0}^{ori} \cdot \lambda_c^{ori} \cdot \prod_{-1 \le p \le 1\ p \ne 0} \lambda_{t,p,n_p}^{ori} & where\ c(t) = III \end{cases}$$

### 2. Mitochondrial de novo mutation dataset

We aimed to estimate the de novo mutation rate for every possible base substitution in the mtDNA. Therefore, we used available datasets of mitochondrial de novo mutations to quantify mutability. We manually compiled a list of mitochondrial de novo mutations from the literature and supplemented these with an in-house dataset (Supplementary Table 2). This included germline de novo mutations, as well as somatic mutations (acquired in tissues and cancer) which have highly similar mutational mechanism and signature to germline mutations in the mtDNA[2,6].

**Supplementary Table 2: Overview of the de novo mutation datasets used.** The reference, associated file used, and any other relevant information is provided.

| Dataset | Source and reference | Notes |
|---|---|---|
| Dataset 1 | Germline[4] | Marked as de novo in Data S1. |
| Dataset 2 | Germline and somatic tissue[7] | Germline de novo is marked as 'child', and somatic as 'somatic-gain', in Table S3. |
| Dataset 3 | Germline and somatic tissue[8] | Table S3 (germline) and Table S4 (somatic tissue) was obtained from the Appendix file, and converted to txt file format. Dataset S1 was used to determine the major and minor allele at each position in both tables. |
| Dataset 4 | Germline[9] | De novos were manually identified from Dataset S1. |
| Dataset 5 | Germline* | In-house dataset, described in section 2.1. |
| Dataset 6 | Somatic tissue[6] | Provided in Table S3. Inferred as aligned to the Yoruban reference sequence, so converted to the rCRS. |
| Dataset 7 | Somatic cancer[10] | In the file retrieved from https://ibl.mdanderson.org/tcma/download/TCMA-MutationSNV.tsv.zip |

*This de novo dataset is unpublished.

We first performed checks to ascertain if there were any major differences between the de novo mutations identified from the various sources (germline, somatic tissue, and somatic cancer), as well as from samples with higher de novo counts. We compared the mutation likelihoods produced from each source of de novo mutations, for transitions. This revealed that while the likelihoods estimated by germline and somatic tissue datasets were similar, the likelihoods estimated by somatic cancer mutations for G>A (and to a lesser extent A>G) mutations diverged as the total number of de novo mutations per sample increased (Supplementary Fig. 1). Therefore, we decided to only use somatic cancer mutations that were from samples with only one de novo mutation, since these were similar to germline and somatic tissue datasets, and given we ultimately aimed to estimate the germline de novo rate.

This resulted in a final list of 4216 de novo mutation allele counts, including 3799 within the reference sequence excluding the OriB-OriH region (m.192-16196, including 3421 transitions and 378 transversions), and 417 within the OriB-OriH region (m.16197-191, including 385 transitions and 32 transversions).

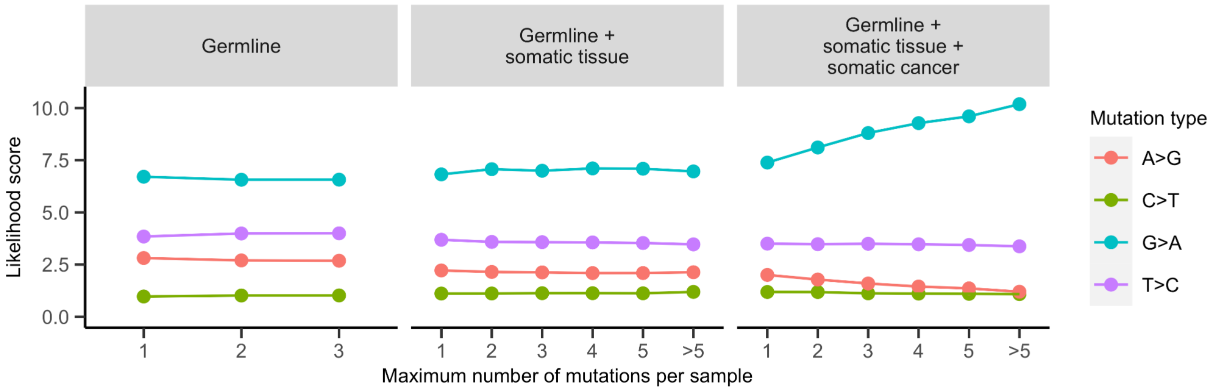

**Supplementary Figure 1: Mutation likelihoods of de novo transitions by source.** The likelihood of G>A mutations increases linearly as the sample de novo mutation count increases.

#### 2.1 In-house de novo mutation dataset

De novo mtDNA mutations identified from Simons Foundation Powering Autism Research for Knowledge (SPARK)[11] study participants were included in the de novo dataset used for mutational model building, obtained as follows. Whole genome sequencing data from SPARK releases 1-3 were retrieved from SFARI Base for probands and their family members. These data are subject to quality control prior to release, including relatedness and contamination checks. Only unaffected siblings with maternal data were used to generate de novo calls, since the mtDNA has been implicated in autism pathogenesis[12]. This left 1690 mother-child pairs for analysis. GATK Mutect2 “mitochondria-mode” was used to call variants, as described previously[13] with the exception that filters “possible_numt” and “mt_many_low_hets" were also applied. All PASS variants in each child at >1% heteroplasmy level that were not a PASS variant with >1% heteroplasmy in their mother were regarded as candidate de novo variants. Filtering was then applied to produce a high-confidence list by excluding any candidate de novos where (1) a variant was called at same position in the mother, including non-PASS calls (i.e. a different PASS variant, or the same variant was non-PASS in the mother), or (2) if the candidate de novo was a SNV and an indel was called at the same or an adjacent position (+1 and -1 bases) in the child (including non-PASS indels). From this a list of 540 de novo mutations were identified, a de novo rate consistent with that reported by Wei et al[4], which were used in the de novo dataset.

**3. Validation of mutation model**

The observed level of neutral variation in a population dataset should correlate with mutation rates, since it is not subject to strong selection. Therefore, we measured the correlation between the observed level of neutral variation in gnomAD and their mutation likelihoods, to confirm the predictive value of our mutational model. We used haplogroup variants from PhyloTree to represent neutral variation given they are passed down a maternal lineage at homoplasmy, supplemented with variants at non-conserved sites to increase representation of different mutation types (i.e. transversions), which we defined as being in the lowest decile of PhyloP scores (equivalent to a score of <-7.3 for reference excluding OriB-OrH and <-2.8 for the latter). This approach enabled us to include neutral variation in protein and RNA genes, and in non-coding loci.

We fit the highly mutable G>A and T>C variants separately, akin to how CpG transitions are handled separately in nuclear models[14], given prior analyses had shown that these mutation types were likely saturated in gnomAD[13]. Neutral variation was assessed in all mtDNA genes as well as in the non-coding sequence excluding OriB-OriH. We observed that the mutation likelihoods were highly correlated with the observed level of neutral variation in each locus in gnomAD, measured as the observed sum maximum heteroplasmy, with Pearson correlation coefficients of R>0.99 (Supplementary Fig. 2a, p-value <2.2e-16), which remained high when restricting to the smaller tRNA genes (Supplementary Fig. 2b). A comparison of correlations using the *cocor* R package[15] showed that the correlation between mutation likelihoods and observed levels (R=1) was significantly stronger than that between mutation likelihoods and locus length (R=0.93, *cocor* p-values <1x10$^{-03}$). Multiple regression analysis confirmed that the association between the mutation likelihoods and observed variation (p<2.2e-16 for G>A/T>C and all other mutations) was much more significant than that between locus length and observed levels (p=0.03 and p=0.134 for G>A/T>C and all other mutations respectively). We validated the non-coding OriB-OriH region separately to the rest of the reference sequence. To enable assessment of correlation in this region, we segmented it into eight non-overlapping blocks of approximately equal size, of similar length to the tRNA genes (70 bp). We observed a high correlation between mutation likelihoods and observed neutral variation within the OriB-OriH region (R>0.9, p<3.1e-05 and p<0.002) (Supplementary Fig. 2c). These data establish the predictive value of the mutational model.

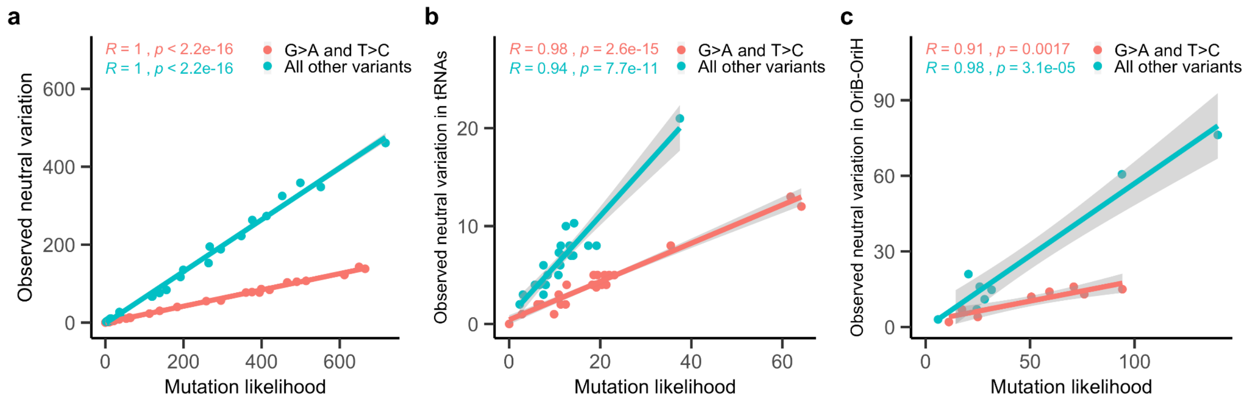

**Supplementary Figure 2: Assessing the predictive value of the mutational models. (a-c)** For each locus **(a)**, tRNA gene **(b)** or equally sized blocks in the OriB-OriH region **(c)** the observed sum maximum heteroplasmy of neutral variants is plotted against the sum of their mutation likelihood scores. The highly mutable G>A and T>C were fit separately (shown in red). Pearson correlation coefficient (R) and its p-value (p) is shown. The line represents the linear model fit, and the gray shading the 95% confidence level around it.

### 4. Assessment of mitochondrial constraint

We assessed mitochondrial constraint in gnomAD as a ratio of observed to expected variation. Nuclear models of constraint applied to gnomAD assess the number of unique variants within a gene or region in the population[14]. Since selection can occur against both the number of variants and their heteroplasmy level in the mitochondria[13], we assess the observed and expected sum maximum heteroplasmy of mtDNA variants in the population. This value represents the maximum heteroplasmy (fraction of alternate to total reads) that the variant is observed at across all individuals in gnomAD. Simulation of germline mtDNA mutation and heteroplasmy drift across generations support the validity of using the mutational model to estimate the expected sum maximum heteroplasmy (see below). Each mtDNA variant in gnomAD v3.1 has a maximum heteroplasmy between 0.1-1.0 since only variants with heteroplasmy level >10% were included in this release; variants not observed are assigned a maximum heteroplasmy of 0.

The observed value for a variant class and or within a gene or region is determined by summing the maximum heteroplasmy of every possible single nucleotide variant (SNV) belonging to this group (e.g. all missense within a protein-coding gene). We calculated the expected sum maximum heteroplasmy by summing the mutation likelihoods of every possible variant within the variant class/region of interest, and then applying the linear equations fit on mutation likelihoods and observed neutral variation in gnomAD; the highly mutable G>A and T>C variants were fit separately, as were variants within the non-coding OriB-OriH region. Variants at six artifact prone sites not called in gnomAD were excluded from observed and expected value calculations (m.301, m.302, m.310, m.316, m.3107, and m.16182)[13]. We applied a clamp to bound the expected value by biological limits, such that it cannot be less than 0 or greater than the maximum possible value. This method can currently only be applied for the assessment of SNVs, and therefore provides the observed:expected ratio for SNVs specifically.

We further provide a 90% confidence interval (CI) around the observed:expected (oe) ratio, adapting the method used for nuclear constraint models applied to gnomAD[14]. For a given pair of observed and expected values, we compute the density of a beta distribution where $x$=proportion possible observed, and parameters $a$=expected value times a varying parameter (ranging between 0-2) + 1, and $b$=maximum possible observed - $a$. The cumulative density function of this function is computed and the value of the varying parameter is extracted at points corresponding to 5% and 95% to indicate the bounds of the CI around the oe ratio. The oe ratio 90% CI upper bound fraction (OEUF) provides a conservative measure of constraint and is used for most analyses.

Nuclear constraint models include corrections for low coverage regions and methylation levels[14]. We note that we do not apply these given the high and even coverage of the mtDNA in gnomAD[13], and lack of robust data on mtDNA methylation[16,17].

**5. Simulation of germline mtDNA mutation and heteroplasmy**

To support the utility of using the mutational model to assess maximum heteroplasmy, we adapted a computational framework developed by Colnaghi et al[18] to simulate germline mtDNA mutation and heteroplasmy drift across generations, and used it to validate a correlation between mutation rates and maximum heteroplasmy for neutral mutations. We applied the parameters drawn from human data by Colnaghi et al, and adapted their approach of modeling mtDNA replication and cell division during oogenesis (development of the female germline) as follows:

(1) Starting from a zygote with $2^{19}$ mtDNA copies (approximately half a million), we first simulated the formation of a primordial germ cell (PGC) through a series of 12 cell divisions without mtDNA replication per Colnaghi et al, with random partition of mtDNA to the daughter cells at each division drawn using a hypergeometric distribution. This results in the formation of a bottleneck, such that the PGC has 128 mtDNA copies (or fewer as specified).

(2) We then modeled the formation of a primary oocyte from the PGC via a series of 18 cell divisions with mtDNA replication per Colnaghi et al, during which constant mtDNA copy number is maintained. This involves first modeling the probability of a mutation during replication, achieved via a binomial distribution for 10 mutation rates between $10^{-9}$-$10^{-7}$ per base pair, and then simulating cell division as above. We also included the possibility of back mutation during replication at the same mutation rate and assumed that the PGC survives the random cell death that occurs during this process.

(3) We then simulated the formation of a mature oocyte from the primary oocyte, via clonal amplification of the mtDNA without cell division until the starting mtDNA copy number of $2^{19}$ is reached. The probability of mutation during replication is modeled as above. We extend this framework across multiple generations by assuming that the heteroplasmy level in the mature

oocyte is equal to the heteroplasmy level in the zygote it forms, providing the starting heteroplasmy level for the next generation.

This process from zygote to mature oocyte was repeated for five generations, for 10,000 maternal lineages. This method has a few simplifying assumptions; we follow the heteroplasmy of only one mutation per lineage, we assume that it is neutral and not subject to selection, and we assume one pool of mtDNA per cell such that we don't consider multiple segregating units. We evaluated the validity of our adaptation by reproducing an analysis reported by Colnaghi et al[18] and Wei et al[4]; the results of the simulation analysis are described in Extended Data Fig. 2.

**6. Regional constraint**

Although motivated by analyses of nuclear regional missense constraint[19], these methods were not applicable to the mtDNA, in part due to utilizing exon boundaries. Therefore we developed a method that compares the missense observed:expected (oe) ratio of all possible regions within a gene to the gene itself, to identify any regions which have a missense oe ratio that is significantly lower than the gene, an approach that is computationally feasible within the mtDNA. We then adapted the approach taken by Davydov et al[20] to reduce these data to discrete intervals of regional missense constraint by applying a greedy algorithm, and calculated the false discovery rate of each using random permutations of the reference sequence. The final method applied is as follows.

First, a list of all possible regions within each protein gene (termed *k*-mers) was compiled, of lengths ranging from a user-defined minimum to the maximum possible (gene length - 1). These *k*-mers spanned codons and *k*-mers 10 codons or larger were assessed (i.e. p.1-10); this minimum was used to enable all *k*-mers to have expected value >10 (Supplementary Figure 3a). Starting from the first codon position, a *k*-mer of the minimum length is drawn (e.g. p.1-10) and the missense oe ratio within it is calculated (where missense is the most severe consequence). The *k*-mer length is then increased by one codon (e.g. p.1-11), and the missense oe ratio within this *k*-mer is calculated. This process is repeated until all possible lengths with that *k*-mer start position are evaluated (e.g. p.1-12, p.1-13, etc), after which the *k*-mer start position is moved by one and the process is repeated (e.g. p.2-12, p.2-13, etc). For each *k*-mer, the probability of observing a missense oe ratio that is less than or equal to the gene's missense oe ratio is calculated by computing the cumulative density function of a beta distribution with parameters a=(expected value * gene oe ratio) + 1 and b=maximum possible value - a. All *k*-mers with p-value >0.01 are discarded; we also discard any remaining *k*-mers with a missense oe ratio 90% CI that overlaps the gene's CI. A greedy algorithm is then applied, such that any *k*-mer which overlaps another *k*-mer with a lower (more significant) p-value is discarded; for overlapping *k*-mers with the same p-value the longest *k*-mer is retained. This produces a list of non-overlapping *k*-mers that are significantly more missense constrained than the gene, which we term candidate areas of regional missense constraint (RMC). This process is summarized in Supplementary Fig. 4.

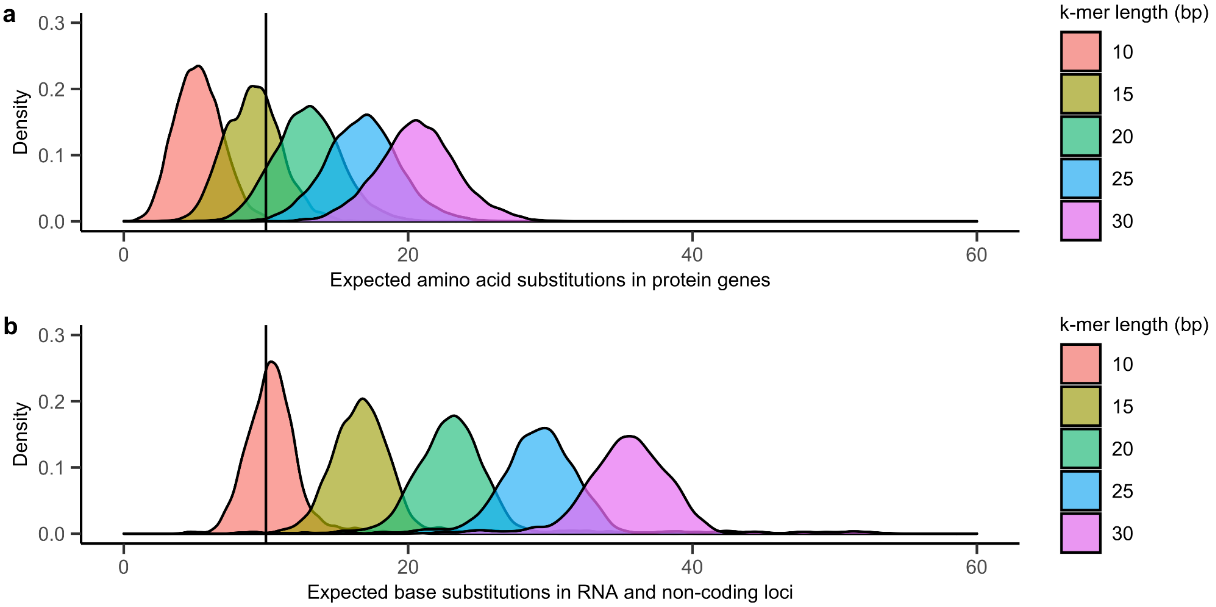

**Supplementary Figure 3: Distribution of expected values for *k*-mer by length. (a-b)** Density plots showing the distribution of expected values across all possible *k*-mers of each length (in bp) for missense variants in protein genes **(a)** or base substitutions within non-protein loci **(b)**. A vertical line marking an expected value of 10 is shown.

We calculated the false discovery rate (FDR) for each candidate area of RMC by assessing random permutations of the reference sequence. Specifically, we generated 1000 permutations of each gene by randomly shuffling the order of the codons, and then applied the process described above to yield a set of non-overlapping significant *k*-mers (if any) which we regard as false positives. We note that the entire gene sequence is permutated, such that any candidate areas of RMC are included, which may contribute false positives. We calculated the FDR of each candidate RMC area by counting how many permutations of the same gene had a false positive result of the same length and ≤ p-value to the candidate, and divide this number by the total number of permutations ($n$=1000). We then filtered out any candidate RMC areas with FDR >0.1. Longer candidates are more likely to have lower FDR, and therefore a filtered candidate may overlap another significant $k$-mer that has a FDR <0.1 but was discarded by the greedy algorithm (i.e. a longer region discarded due to having a higher p-value than the filtered candidate). To recover these, we reapply the greedy algorithm after removing regions with FDR >0.1 from both the real and permuted sequence, and recalculate the FDR for each resultant candidate RMC. The process is repeated if needed until all remaining candidates satisfy the FDR <0.1 threshold, or no candidates remain. We note that only 9 of 39 initial candidate RMC areas did not pass the FDR threshold. This process yields a list of high-confidence RMC areas that are used for all analyses (Supplementary Dataset 2).

Regional constraint in the rRNA genes was calculated using the same process with the following modifications: (1) $k$-mers are measured in bp, (2) all possible base substitutions (i.e. SNVs) are included in observed and expected calculations, (3) $k$-mers 20 bp or larger were assessed (to enable all to have expected value >10 per Supplementary Fig. 3b), and (4) the random permutations of each gene are achieved by shuffling the order of the bases.

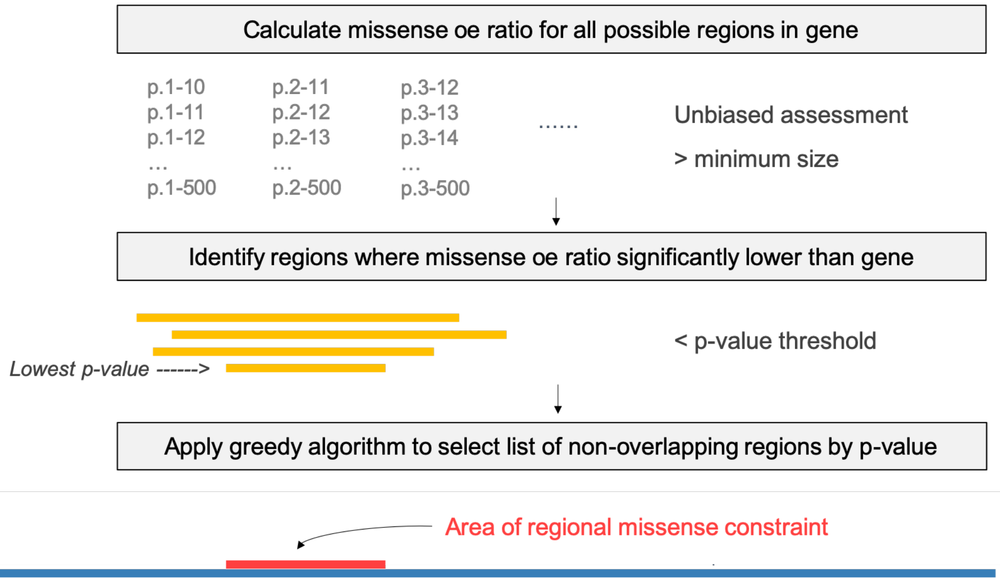

**Supplementary Figure 4: Overview of the regional missense constraint method.** The missense observed:expected (oe) ratio of all possible regions within a gene are assessed to identify any that have a missense oe ratio that is significantly lower than the gene's. A greedy algorithm was then applied to reduce these data to discrete intervals of regional missense constraint by p-value. The same method is applied to random permutations of each gene sequence for estimation of the false discovery rate.

**7. Local constraint score**

We were interested to identify the most constrained sites in the entire human mtDNA. To do this, we employed a sliding window approach for capturing the strength of selection against base or amino acid substitutions at and around each base position, which we term local constraint. From an initial start position of m.1, a window of length *k* is drawn and the observed:expected (oe) ratio and 90% confidence interval (CI) of substitutions within the window is calculated. The window start position is then moved by 1 bp, and the oe ratio and CI of the next window of length *k* is calculated. This process is repeated until all possible overlapping windows of length *k* in the mtDNA are evaluated; windows which overlap two different loci are therefore included. For positions in non-coding and RNA loci all possible base substitutions (i.e. SNVs) are included in calculations, while for positions in protein genes only missense variants are included to restrict assessment to amino acid substitutions (where missense is the most severe consequence). We used a uniform window length *k* of 30 bp, which for proteins captures 10 amino acids; this length was used to enable all *k*-mers to have an expected value >10 (Supplementary Fig. 3).

The local constraint of each base position in the mtDNA was then measured by taking the mean oe ratio CI upper bound fraction (OEUF) of all *k* windows that overlap the position. This approach therefore incorporates data from neighboring bases, such that bases in the closest proximity will contribute more of the signal than bases further away (Supplementary Fig. 5a). These mean OEUF values were then percentile ranked, such that each position was assigned a score between 0 and 1, which we term the mitochondrial local constraint (MLC) score (Supplementary Fig. 5b). The higher the score, the more locally constrained the position is. The score for every base position is provided in Supplementary Dataset 6. Since this score is a measure of local intolerance to missense variants specifically in proteins, we also assigned scores to non-missense variants in protein genes to enable application to all possible SNVs. The MLC scores for variants were obtained as follows: non-coding, RNA and missense variants were assigned their positional score, synonymous variants were assigned a score of 0, stop gain were assigned a score of 1, and start and stop lost variants were assigned a score of 0.70. These assigned values for non-missense variants in protein genes are based on the OEUF of these functional variant classes (synonymous OEUF=1.01, stop gain OEUF=0.015, start lost OEUF=0.35, stop lost OEUF=0.34) and their placement within the spectrum of mean OEUF values (Supplementary Fig. 5b). MLC scores for every possible mtDNA SNV are provided in Supplementary Dataset 7.

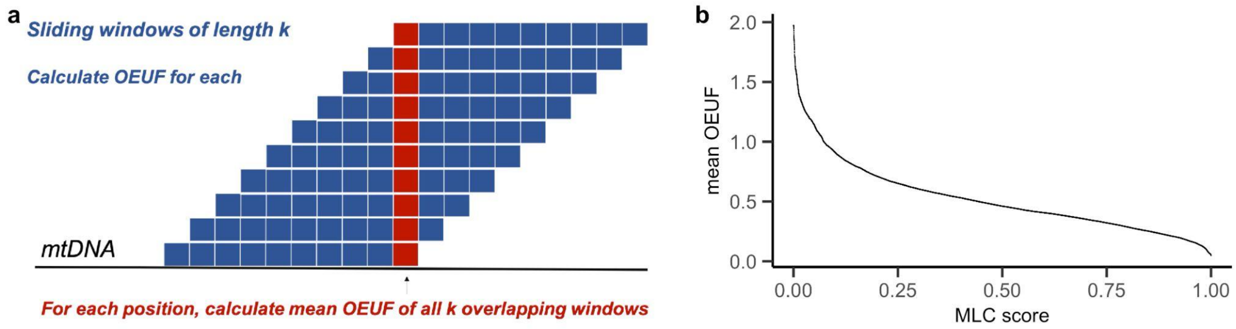

**Supplementary Figure 5: Overview of the mitochondrial local constraint (MLC) score. (a)** Schematic overview of the method used to generate the MLC score. A sliding window of uniform length *k* is moved across the mtDNA at +1 base pair intervals. The observed:expected (oe) ratio of amino acid or base substitutions, and its 90% confidence interval (CI), was calculated for each window. For positions in protein genes only missense variants were included in calculations, while for all other positions all base substitutions are used. The mean oe ratio CI upper bound fraction (OEUF) of all *k* overlapping windows is computed for each position, and then percentile ranked to produce the MLC score. Note non-missense variants in protein genes are handled separately. **(b)** Relationship between the mean OEUF and MLC score.

### Supplementary Discussion

#### 1. Extended discussion on simulation of germline mtDNA mutation and heteroplasmy

We adapted a computational framework developed by Colnaghi et al[18] to simulate germline mtDNA mutation and heteroplasmy drift across generations, and used it to validate a correlation between mutation rates and maximum heteroplasmy for neutral mutations. The validity of our adaptation was supported by reproducing the results of an analysis reported by Colnaghi et al[18], which assessed the heteroplasmy distribution for five different bottleneck sizes in offspring who inherited a mutation with heteroplasmy 0.1 (Extended Data Fig. 2a). Changes in heteroplasmy levels between generations was also similar to real-world data reported by Wei et al from ~1500 mother-offspring pairs[4], further supporting the utility of our model (Extended Data Fig. 2b). We then evaluated maximum heteroplasmy across five generations for 10 different mutation rates, using a starting heteroplasmy of 0 for generation 0 (Extended Data Fig. 2c). We used bootstrapping to assess the maximum heteroplasmy distribution at generation five for each mutation rate tested. This showed an approximately linear correlation between the mutation rate and the median maximum heteroplasmy for neutral mutations, such that higher mutation rates had higher maximum heteroplasmy, supporting our application of the mutational model for the estimation of maximum heteroplasmy values (Extended Data Fig. 2d). This is consistent with conclusions drawn by Ju et al, who found that the expected number of neutral mitochondrial mutations drifting to homoplasmy increases linearly with mutation rate in tumor cells[2].

#### 2. Extended discussion on regional missense constraint within protein genes

Manual inspection of regional missense constraint within protein structures revealed multiple examples of their topological clustering in 3D space. This includes for complex I subunits MT-ND1 (Fig. 2b) and MT-ND6 (Extended Data Fig. 5e), and complex III subunit MT-CYB (Extended Data Fig. 5d). Review of the literature further revealed that several of these topological clusters of regional constraint were notable for encoding residues of functional significance, as were other areas of regional constraint. For example MT-ND6 plays a key role in the transition between the ‘open’ and ‘closed’ states of complex I, which is critical for its catalytic activity[21]. Notably, the areas of regional constraint within this protein encode or are located in close proximity to residues critical for this function including within the π-bulge and interface-forming loop[21] (Extended Data

Fig. 5e). Complex III is responsible for electron transfer from ubiquinol to cytochrome c as part of oxidative phosphorylation, and it is MT-CYB that catalyzes this process. The four areas of regional constraint in MT-CYB cluster to the sites of the heme molecules and nearby quinone oxidation and reduction sites, which are critical for this function[22,23] (Extended Data Fig. 5d). Residues in the ef loop (p.252-268), PEWY sequence (p.269-283) and cd1 helix (p.136-152) which are also critical for electron transfer[23] also overlap regional constraint (Supplementary Dataset 2). The one area of regional constraint identified in complex V subunit MT-ATP6 is also notable for being located at the interface with the c-ring, within an area critical for proton translation and ATP synthesis[24]. Collectively, these observations support that regional missense constraint identifies areas critical for protein function.

#### 3. Extended discussion of regional constraint within rRNA genes

There are only a handful of confirmed pathogenic variants in the mitochondrial rRNA genes, and therefore the strong negative selection observed against rRNA variation is particularly striking. However, this can be reconciled with the fact that the rRNAs have an essential role in the mitochondrial ribosome (mitoribosome) and translation. In support of this, the most regionally constrained interval in a rRNA gene (m.3028-3081 in MT-RNR2) encodes the A-loop which interacts with the 3' end of an aminoacylated tRNA in the A-site[25]. This the first binding site for a tRNA in the mitoribosome during protein synthesis, and therefore likely one of the most functionally important areas of the rRNA. This area of regional constraint has an OEUF value of 0.07 (Supplementary Dataset 2), which is similar to pLoF OEUF gene values, supporting that many variants in this region are deleterious. Indeed, no homoplasmic variants were observed in this region in the gnomAD and HelixMTdb population databases collectively comprising data from up to ~250,000 individuals[13,26]. Only five variants in this region were reported in MITOMAP[27]; these were each reported in only one sample, and manual inspection showed that they were enriched in sample types with increased tolerance to deleterious variation and or lower quality (m.3033T>C in sample with GenBank ID LS998724.1 deposited by a Department of Pediatrics Genetic Diagnostic Laboratory, m.3058T>C in KJ735669.1 from cells, and m.3077C>T in MH043584.1 from ancient DNA)[27]. To contextualize this observation, a similarly sized region in MT-RNR2 (m.2220-2270) has an OEUF value of 0.80 and >40 homoplasmic variants reported in gnomAD and HelixMTdb databases[13,26]; MITOMAP also has >40 variants in this region[27]. This example showcases a functionally critical rRNA region that may harbor overlooked deleterious variation, and supports our evidence of constraint within rRNAs, particularly within MT-RNR2 which has not had any *bona fide* pathogenic variants reported.

#### 4. Extended discussion of local constraint score across the non-coding control region

The signal of constraint across the artificial chromosome break at m.16569-1 is unexpected (Extended Data Fig. 7a). Pipelines which use the linearized mtDNA reference sequence can be vulnerable to undercalling variants around the break; however the Mutect2 pipeline utilized for gnomAD used a shifted alignment to specifically avoid this issue[13]. The artificial chromosome break also lies within the OriB-OriH region, which is known to have a different mutational signature[2,10] and was handled separately by our model (see Supplementary Methods). Future work is required to clarify if mutational properties and or variant calling challenges unique to this region could be contributing to this signal of constraint. It is notable however that all of the other signals of constraint within the non-coding control region overlap annotated or unannotated regions previously implicated as functionally important (Extended Data Fig. 7a). Two of these regions were only recently established to have an essential role in mitochondrial transcription, including the conserved sequence block 3[28] and the second light strand promoter[29], supporting that studies into the functional role of the region around the artificial chromosome break may also be warranted. Indeed, this region lies downstream of the origin of heavy-strand replication and nearby to the 5′ ends of 7S DNA, a linear strand of DNA which forms a triple-stranded structure with the D-loop which is hypothesized to have functional significance[30].

#### 5. Extended discussion on the local constraint scores of disease-associated variation

The mitochondrial local constraint (MLC) scores of disease-associated variation shows enrichment of pathogenic variants in the higher score quartiles (Fig. 5c). However, some pathogenic variants have low scores (Extended Data Fig. 8a-d). There are several possible explanations for this. Firstly, the MLC score incorporates data from neighboring bases to measure local tolerance to variation, such that the bases in the closest proximity will contribute more of the signal. Therefore, deleterious variation could be assigned low scores if they reflect the mutation of a single critical residue or base in a neighborhood that is mostly tolerant of variation. This also means that positions with the highest scores are more likely to lie in sites where multiple neighboring residues/bases are critical for function. Accordingly, a benign variant could have a high score due to being in a neighborhood which is largely intolerant of base/amino acid substitution. Secondly, for missense and RNA/non-coding variants the same score is assigned for different variants at the same position, which therefore doesn't take the type of amino acid or base change into account. It's possible that a specific substitution has deleterious effects where others in the local area are tolerated. Thirdly, the heteroplasmy level that the deleterious variation is tolerated at could affect the measure of selection, such that a neighborhood that can tolerate deleterious variation at homoplasmy will have a lower score than those where strong selection occurs against lower heteroplasmy variants (e.g. where homoplasmy may not be compatible with life). This is supported by the observation that confirmed disease-associated variants reported to cause disease at homoplasmy have an increased density of scores <0.75 compared to those with a disease association at heteroplasmy only, per their status in MITOMAP (Extended Data Fig. 8c); a trend which was strengthened following manual curation of heteroplasmy and homoplasmy status the literature (Methods) (Extended Data Fig. 8d). The MLC score is therefore positioned to be particularly useful for capturing the effect of deleterious heteroplasmic variation.

**6. Extended discussion of the association between the MSS and blood cell counts**

Application of the mitochondrial local constraint score sum (MSS) metric supports that deleterious mtDNA variation likely plays a causal role in platelet counts, and a non-causal role in neutrophil counts. This is in line with mtDNA having a critical role in platelets due to their lack of nuclear DNA[31], and with the diminished role of mitochondria in energy generation and other cellular functions in neutrophils[32]. The results showed a positive correlation between heteroplasmy count and neutrophil count, as well as between the MSS and platelet count (Fig. 5d). The latter data point suggests that an increased burden of deleterious mtDNA variation leads to increased platelet count. While initially counterintuitive, this may reflect a possible compensatory mechanism. Support for this in the literature includes data showing that an increase in mitochondrial reactive oxygen species (a marker and consequence of mitochondrial dysfunction) can lead to platelet biogenesis[33], and evidence that platelet apoptosis can paradoxically lead to a higher proportion of younger platelets[34]. Further work is required to elucidate the possible mechanisms underlying these data.

### Supplementary Datasets

**Supplementary Dataset 1: Constraint metrics for human mtDNA genes.** Gene symbol and coordinates, and the observed and expected sum maximum heteroplasmy of SNVs in the gene in gnomAD v3.1, observed:expected ratio, and lower and upper bounds of the 90% confidence interval (CI) around the ratio are listed, including for each major class of protein variation (synonymous, missense, stop gain) for protein genes.

**Supplementary Dataset 2: Intervals of regional constraint in mitochondrial proteins and rRNA genes.** Gene symbol and coordinates of regional constraint, and the observed and expected sum maximum heteroplasmy of SNVs in the region in gnomAD v3.1, observed:expected ratio, and lower and upper bounds of the 90% confidence interval (CI) around the ratio are listed. Note this represents regional missense constraint for protein genes, and regional constraint for RNA variants in rRNA genes.

**Supplementary Dataset 3: Curated missense variants identified in cases suspected to have a primary mitochondrial disorder.** Curation and classification were performed by the Victorian Clinical Genetics Service, and variant position, reference and alternate alleles, and their classification group are listed. These data are shown in Fig. 2d.

**Supplementary Dataset 4: Constraint metrics for each position within the tRNA secondary structure.** The tRNA secondary structure position number, and the observed and expected sum maximum heteroplasmy of SNVs at the position in gnomAD v3.1, observed:expected ratio, and lower and upper bounds of the 90% confidence interval (CI) around the ratio are listed.

**Supplementary Dataset 5: Constraint metrics for annotated non-coding elements.** Locus name, description and coordinates, and the observed and expected sum maximum heteroplasmy of SNVs in the element in gnomAD v3.1, observed:expected ratio, and lower and upper bounds of the 90% confidence interval (CI) around the ratio are listed.

**Supplementary Dataset 6: Mitochondrial local constraint (MLC) base scores for every base position.** The mtDNA position and its position score is listed. Note this score measures missense tolerance specifically for positions in protein genes (Methods).

**Supplementary Dataset 7: Mitochondrial local constraint (MLC) variant scores for every single nucleotide variant.** The variant position, reference and alternate alleles, consequence and MLC score are listed. Note that non-missense variants in proteins were assigned scores separately (Methods).

**Supplementary Dataset 8: Confirmed pathogenic variants with a curated disease plasmy status.** Disease-associated variants with a 'confirmed' status and plasmy status of '-/+' in MITOMAP had their plasmy status in the literature manually reviewed. The variant, its gene symbol and MITOMAP plasmy status, status after curation, and PMIDs of publications used for the curated status are listed. Note only missense and RNA variants are included. These data are shown in Extended Data Fig. 8d.

### Supplementary Videos

**Supplementary Video: Mitochondrial local constraint across the 16S rRNA encoded by MT-RNR2.** The video shows the mitoribosome, a complex of proteins (blue) and the small 12S (pink) and large 16S (purple) rRNAs, which serves as the site of mitochondrial translation. The mRNA (bright green) and tRNAs occupying the A/P-site (yellow) and P/E-site (orange) are also shown. The mitochondrial local constraint scores are then displayed across the 16S rRNA encoded by MT-RNR2 using a red-white-blue gradient. Dark red indicates highly constrained sites with scores close to 1, white scores around 0.5, and dark blue scores approaching 0.
